# Neutral genomic signatures of host-parasite coevolution

**DOI:** 10.1101/588202

**Authors:** Daniel Živković, Sona John, Mélissa Verin, Wolfgang Stephan, Aurélien Tellier

**Affiliations:** Section of Population Genetics, Technical University of Munich, 85354 Freising, Germany; Department of Mathematics and Statistics, Queen’s University, Ontario K7L 3N6, Canada; Leibniz Institute for Evolution and Biodiversity Science, 10115 Berlin, Germany

**Keywords:** population genomics, epidemiological model, host-parasite coevolution, SI model, population demographic history, population dynamics

## Abstract

Coevolution is a selective process of reciprocal adaptation between antagonistic or mutualistic symbionts and their host. Classic population genetics theory predicts the signatures of selection at the interacting loci but not the neutral genome-wide polymorphism patterns. We here build a coevolutionary model with cyclic changes in the host and parasite population sizes. Using an analytical framework, we investigate if and when these population size changes can be observed in the neutral site frequency spectrum of the host and parasite full genome data. We show that polymorphism data sampled over time can capture the changes in the population size of the parasite but not of the host because genetic drift and mutations occur on different time scales in the coevolving species. This is due to the small parasite population size at the onset of the coevolutionary history subsequently undergoing a series of strong bottlenecks. We also show that tracking coevolutionary cycles is more likely for a small amount of parasite per host and for multiple parasite generations per host generation. Our results demonstrate that time sampling of host and parasite full genome data are crucial to infer the co-demographic history of interacting species.

## Introduction

Host-parasite antagonistic interactions are a role model for observing and studying rapid evolutionary change as well as feedbacks between ecological and evolutionary forces and time scales. Coevolution, defined here as the reciprocal adaptation of hosts and their parasites, typically generates significant phenotypic and genetic diversity for host resistance and for parasite infectivity and virulence. Such changes in the genetic composition of the interacting species at the key underpinning loci, drive, and are driven by, short-term epidemiological (ecological) dynamics. To develop infectious disease epidemiology as a predictive science, there is thus a need to understand the synergy of fast evolution and within and between populations disease dynamics [1], the so-called eco-evolutionary feedbacks [2].

Coevolution as determined by changes in allele frequencies over time at the interacting genes, is observable as coevolutionary cycles driven by negative indirect frequency-dependent selection [3, 4]. Theory predicts that a continuum of dynamics of allele frequency cycles occurs, characterized by their stability, period and amplitude, and ranging between two extremes: the arms race and the trench warfare dynamics. The arms race is defined as the recurrent fixation of alleles at these major loci in host and parasite populations [5, 6], while trench warfare maintains cycling over a long period of time [7] (also called the Red Queen dynamics [6], or Fluctuation Selection Dynamics [2]). The transition between these types of dynamics depends on the occurrence and strength of negative direct frequency-dependent selection [4], which stabilizes cycles and is generated by several host and parasite life history traits (reviewed in [8]).

Theory also predicts that the coevolutionary dynamics, either by arms race or trench warfare, can be observed in the patterns of polymorphism at these loci, namely in the frequencies of Single Nucleotide Polymorphisms (SNPs). The arms race is expected to generate recurrent selective sweeps, while trench warfare generates balancing selection. These predictions form the basis of scans for genes under coevolution in host or parasite genomes relying on the prevalent perception that natural selection acts only at few loci, while neutral forces, such as demographic histories, affect the whole genome. Detecting genes under coevolution entails therefore to disentangle the signatures of arms race or trench warfare from the polymorphism patterns observed in genome-wide data.

Besides the allelic coevolutionary cycles at the host and parasite interacting loci, size fluctuations of host and parasite populations are also predicted to occur. These changes in population size over time are induced by reciprocal selection among the antagonists and are an inherent property of host-parasite coevolution under epidemiological (or Lotka-Volterra) dynamics (such as the Susceptible-Infected or Susceptible-Infected-Recovered models, [9]). In a more complex coevolutionary system with several host and parasite genotypes being present at major genes of interaction, cycles of coevolution do occur, thereby generating a fluctuation of the numbers of hosts and parasites over time [9] as an epidemiological feedback [2]. Several episodes of coevolution proceed with increasing and decreasing disease prevalence depending on the cycling of resistance and infectivity alleles. The epidemiological feedback generates negative direct frequency-dependent selection, thus stabilizing the frequencies of alleles and maintaining long-term diversity at the interacting loci [10]. Coevolutionary models based on Lotka-Volterra dynamics have similar characteristics [11−14]. Population size changes due to coevolution should affect the whole genome polymorphism of both antagonistic species, an effect which we term as the co-demographic history. When studying host and parasite polymorphism data, two sources of demographic variation generating genetic drift can therefore be defined: 1) the population or species demographic history (*e.g.* colonisation of new habitats or recolonisation), and 2) the co-demographic history due to co-evolutionary and epidemiological dynamics. Both types of demographic events affect the ability to detect genes under coevolution using scans for arms race or trench warfare signatures. Moreover, there is currently no theoretical prediction regarding the signature of co-demographic history on genome-wide polymorphism in hosts and parasites.

Our aim in this study is to propose the first model of neutral polymorphism generated by the co-demographic history of host and parasite populations. First, we establish an epidemiological model describing changes in the numbers of healthy and infected hosts over time focusing on biallelic gene-for-gene and matching-allele infections and initially assigning one parasite per host. Second, we utilize an analytical result [15] for the neutral site frequency spectrum (SFS) under arbitrary deterministic population size changes and apply it to the host and parasite populations. We show that these population size changes can be quite drastic in the parasite and occur on a time scale slow enough to leave a corresponding signature in the SFS over time. Conversely, changes in the host size are barely detected in the polymorphism data. Finally, since such recurrent bottlenecks in parasites cause a reduced amount of polymorphism, we further discuss the impacts of multiple parasites per host and multiple parasite generations per host generation.

## Theoretical and computational framework

### Modeling a single coevolving locus

We have a haploid one locus model with *A* alleles in the host and in the parasite. The infection matrix is given by 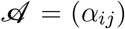 with 1 ≤ *i, j* ≤ *A*. The entries *α_ij_* give the probability that once encountered hosts of genotype i are infected by parasites of genotype *j*. Examples of simple infection matrices for two alleles are given in Table 1.

**Table 1.** Infection matrices for four coevolution models. The infection matrices determine the outcome of the interaction between host genotypes (rows) and parasite genotypes (columns). To keep the illustration simple, the rates *α_ij_* are either chosen as one for infection or as zero for resistance. Matching-allele and gene-for-gene models are shown as well as their inverse versions.

| matching-allele | inverse matching-allele | gene-for-gene | inverse gene-for-gene |
| --- | --- | --- | --- |
| $\begin{pmatrix} 1 & 0 \\ 0 & 1 \end{pmatrix}$ | $\begin{pmatrix} 0 & 1 \\ 1 & 0 \end{pmatrix}$ | $\begin{pmatrix} 0 & 1 \\ 1 & 1 \end{pmatrix}$ | $\begin{pmatrix} 1 & 0 \\ 0 & 0 \end{pmatrix}$ |

In analogy to [16] the changes of host and parasite allele frequencies over time are determined by the following coupled differential equations:

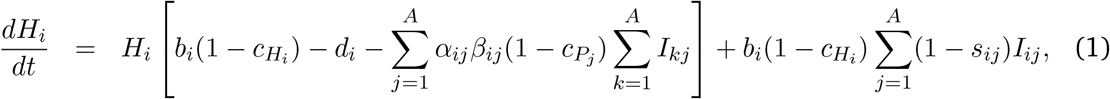

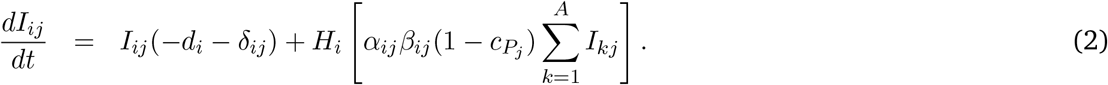

In Equations (1) and (2), *H_i_* is the number of healthy (*i.e.* non-infected) individuals of genotype *i*, and *I_ij_* denotes the number of host genotype *i* infected by parasite genotype *j*. *b_i_* and *d_i_* are the birth and natural death rates (i.e. independent of the disease) of host genotype *i*, respectively, and *δ_ij_* is the disease induced death rate caused by pathogen *j* on host genotype *i* (*i.e.* the effect of pathogen on host mortality [16]). *β_ij_* is the disease transmission rate between a parasite of genotype *j* and a host of genotype *i*. *c_H_i__* and *c_P_j__* are the costs for the hosts and the parasites of carrying genotype *i* and *j*, respectively, *s_ij_* is the decrease of reproductive fitness of host genotype *i* due to an infection of parasite *j*, *i.e.* the effect of pathogen on host fecundity. Due to the large number of parameters in our epidemiological model, we investigate a simplified version with two alleles (A = 2) in the host and in the parasite except for the evaluation of the host effective population size over time (SI 1) and the reproduction ratios (SI 2), which can be computed for arbitrary *A*. Due to the purely deterministic setting, an allele is considered as lost as soon as its count takes a value below one. We only allow rates and costs to differ among the genotypes in SI 1, SI 2, and for the calculation of the fixed points (SI 3) of the dynamical system (1) and (2). Otherwise, we assume *b_i_ = b, d_i_ = d, δ_ij_ = δ, β_ij_ = β* and *s_ij_ = s*. The entries *α_ij_* of the infection matrix are either zero or one depending on the considered model (see Table 1). While we evaluate the fixed points (SI 3) for all four coevolutionary models, only the matching-allele (MA) and the gene-for-gene (GFG) model are investigated in further detail, since the inverse matching-allele (iMA) model is symmetric (and therefore equivalent) to the MA model for *A* = 2 and the inverse gene-for-gene (iGFG) model is restricted in its behavior compared to the GFG model.

For the MA model symmetric costs are chosen and (except for SI 1−SI 3) assumed to be independent of the host (*c_H_i__ = c_H_*) and the parasite (*c_Pj_ = c_P_*) genotype. For the GFG model costs are chosen asymmetrically as follows (except for the choice of arbitrary costs in SI 1−SI 3). Since *H*_1_ defends itself successfully against *P*_1_, and *P*_2_ can infect both host genotypes, only *H*_1_ and *P*_2_ are assumed to carry realistically small costs *c*_*H*_1__ and *c*_*P*_2__ (*c*_*H*_2__ = *c*_*P*_1__ = 0, see [4]).

The total number of hosts of genotype *i* is given by 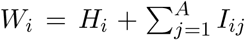. The number *P*_j_ of parasites of genotype *j* is only implicitly given in (1) and (2) by the number of infected individuals; assuming one parasite per infected genotype we have 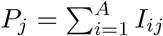.

The change in the effective population size of the host over time 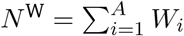 is obtained by numerically solving (1) and (2). The respective differential equation and a condition for obtaining a constant population size are given in SI 1. The effective population size of the parasite is obtained as 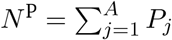 (as we assume first one parasite per host).

### Assessing the effect of the coevolving locus on neutral polymorphism patterns

To evaluate the impact of host-parasite interactions on neutrally evolving and genome-wide distributed SNPs over time for interesting cases (as determined by the stability analysis), we utilize the SFS (SI 4), which is the distribution *f_n,i_* (*t*) of the number of times *i* a mutant allele is observed across sites in a sample of *n* DNA sequences at time *t*. While *f_n,i_* (*t*) takes allele counts in absolute numbers, its relative version 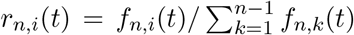 is normalized by the total number of segregating sites. For this purpose, the deterministic trajectories of time-varying host and parasite population sizes are employed into the analytical result [15] for the neutral SFS. This application requires an appropriate scaling: we define a relative population size function *ρ*(*t*) as the ratio of the population size *N*(*t*) at time *t* scaled by the reference population size *N*_ref_ at the time of the infection, which initiates the coevolutionary history, *i.e. ρ*(*t*) = *N*(*t*)/*N*_ref_, and denote the population mutation rate as *θ* = 2 *N*_ref_ *μ*. We compute the changes in frequencies of neutral alleles generated by the co-demographic scenario in terms of the SFS (also in relative numbers) and the average number of pairwise differences 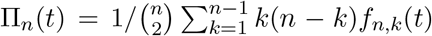 (SI 4) for *n* = 20 hosts and parasites. Our forward approach is adequately suited for the analysis of time-series data in contrast to the corresponding coalescent result for the neutral allelic spectrum [17, 18], where only a single time point can be immediately evaluated for a given demographic history.

### Key characteristics of our model

Our model presents three key features to keep in mind when reading the results below. The first key aspect of our eco-evolutionary framework is that changes in the population size are a direct consequence of the dynamics of the model, driven by a single locus underpinning coevolution, and not assumed to follow an arbitrarily chosen function of time as in the majority of the population genetics literature.

The second crucial point is the definition of the host and parasite time scales of evolution as determined by the generation times and the population mutation rates of the antagonistic species [19]. If viral, bacterial or fungal parasites often have higher mutation rates than their hosts, their effective population size may not always be larger than that of the hosts and at the onset of an epidemics. The reference population size *N*_ref_ at the onset of an epidemics is important because 1) it sets up the initial available diversity, and 2) it defines the time scale for genetic drift in host and parasite and the timing of new neutral mutations occurring with rate *θ*. In our population genetics setting, time is scaled in units of *N*_ref_ generations, whereas the host-parasite model specified in Equations (1) and (2) runs on arbitrary continuous time reflecting calendar time (in weeks, months or years). If calendar time is equivalent for both species, the scaled time based on *N*_ref_ defines the changes occurring in the observed polymorphism over time. We exemplify the difference in time scale and its influence on polymorphism data by a simplified bottleneck model. Two populations with different initial population sizes *N*_ref_ experience a size change of the same magnitude and for the same number of generations on calendar time scale (bottom x-axes) but for different rescaled time with respect to *N*_ref_ (top x-axes of Figure 1, a and c). Consequently, a size change of the same magnitude but based on two different initial population sizes *N*_ref_ affects the SFS similarly regarding the course of time but with very different strength and detectability (y-axes of Figure 1, b and d). Note that the number of singletons scaled by *θ* is used as a representation of the SFS for illustrative purposes in this and several subsequent examples because these show the most pronounced signals of all allelic classes, but results are similar using Π_*n*_(*t*).

**Figure 1.**
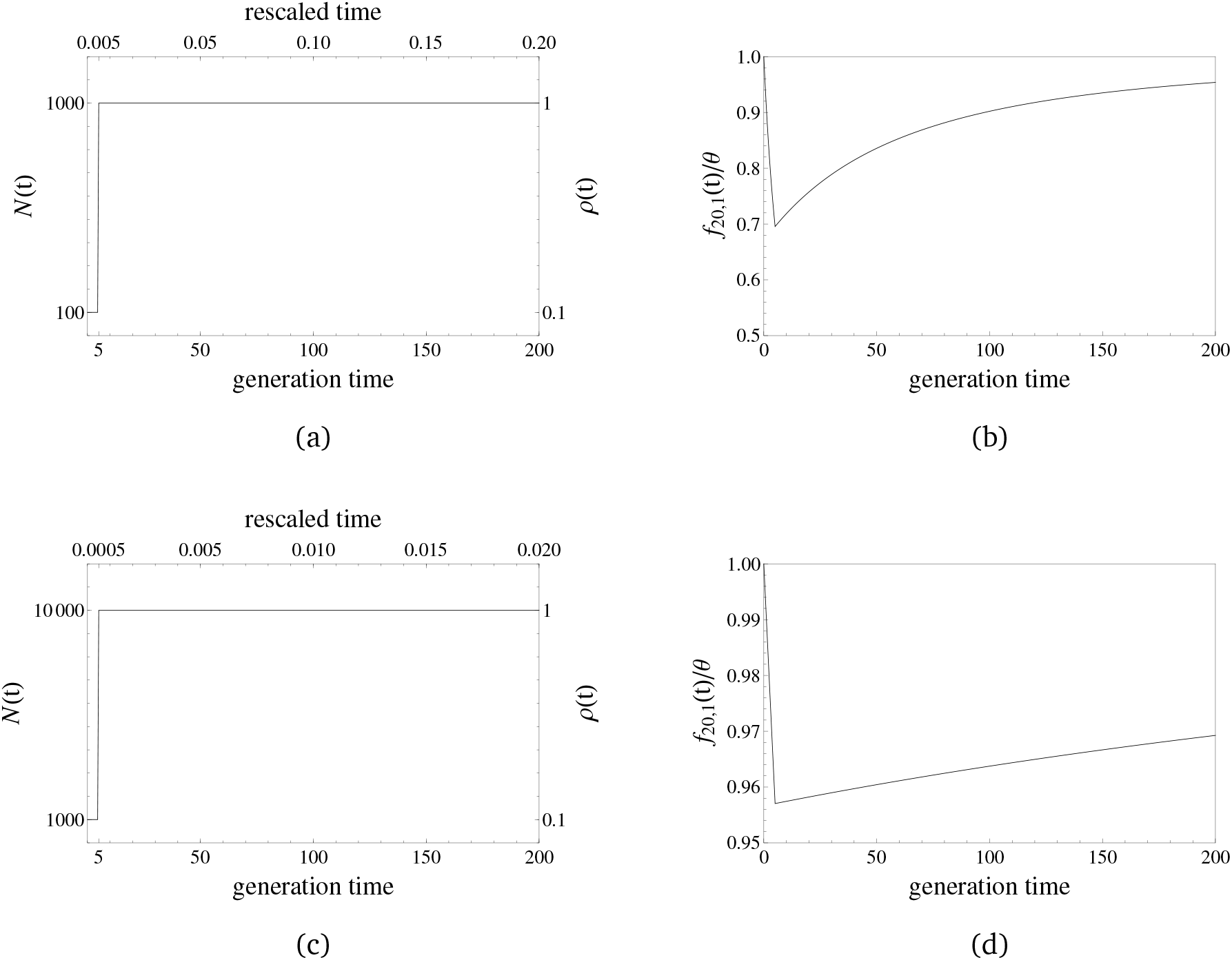
A simple bottleneck model with an initial population size of 1000 (a) and 10000 (c) is considered, where the population size drops instantaneously to one tenth of its original size at time zero for 5 generations before recovering again to its original size. Original time in generations is opposed to time scaled in units of the initial population size on the x-axes and the population size changes are given in absolute and relative numbers on the y-axes. The absolute number of singletons scaled by *θ*, *f*_20,1_(*t*)/*θ*, are illustrated for the respective cases in (b) and (d). The bottleneck is more apparent in (b) than in (d), where the changes of *f*_20,1_(*t*)/*θ* stay within a five percent margin. This is due to slower rescaled time in the first scenario (a), where the population size drop affects polymorphisms prolongedly.

The third important feature of our model is the underlying assumption regarding genetic drift. The theoretical result for our SFS computations (Equation 33 in [15], see also SI 4) is based on a continuous time approximation of the Wright-Fisher model assuming descendants picking their parents at random from non-overlapping discrete generations. However, in our model, there is an overlap of generations in the host and parasite populations to allow the transmission of disease. A higher overlap of generations occurs for lower values of the death rate *d*. In order to apply the theoretical results, we therefore focus specifically on scenarios where the death rate *d* is close to the birth rate *b* = 1, so that most of the population is replaced within a ‘generation’ as defined by 1/*d*. In addition, we build a simulation method that incorporates the overlap of generations in hosts and pathogens during the drawing of the next generation’s allele frequencies.

### Modeling overlapping generations

A simulation protocol was designed that accounts for the effect of overlapping generations in the stochastic sampling of the host and parasite SFS. The differential equations (1) and (2) are discretized by choosing a sufficiently small value for Δ*t* (its infinitesimal version being *dt*) ensuring that the coevolutionary dynamics match the numerical evaluations from *Mathematica*. At every discrete generation *τ*, the current population size *N_τ_* consists of *N_o_*(*τ*) = (1 − *d*) *N*(*τ* − 1) individuals that did not die and therefore overlap and *N_b_* = *N*(*τ*) − *N_o_*(*τ*) newborns (*i.e.* newly infected hosts in case of the parasite). As before, we assume that the SFS of both populations is in equilibrium at start of the infection, *i.e.* the population SFS at time zero is given by *f_x_*(0) = *θ/x* with *θ* = 2 *N*_ref_ *μ* and *μ* being the per generation mutation rate of an entire genome. The SFS of each population is recursively evaluated as follows. For a fixed generation *τ*_0_, the *N*(*τ*_0_) alleles are sampled from the pool of size *N*(*τ*_0_ − 1) and according to the allele frequencies at generation *τ*_0_ − 1. The newborns and the overlap fraction are, respectively, obtained via sampling with and without replacement. New mutants arising only in newborns and as a single copy at a previously monomorphic site are obtained per generation as a Poisson random variable with mean 2*N_b_ μ* and added to the singleton class of the SFS. Once the population SFS *f_x_*(*t*) is computed for a given time interval, its sample version *f_n,i_*(*t*) is readily obtained via binomial sampling. Note that the number of novel mutations and therefore the amount of polymorphism over time is reduced by definition in this model with overlap compared to the Wright-Fisher model, where all individuals are newborns when descendants replace their parental generation.

## Results

### Effect of the various parameters on the dynamical system

We are only interested in situations where at least one host and a parasite genotype survive and both populations coexist. Therefore, we derive first the criteria for the disease to spread in the population via the reproduction ratios (SI 2). Then, we scan the parameter space of our epidemiological model to determine the behavior of the coevolutionary dynamics and the speed of cycling. We thus eliminate situations where the disease spreads but the host and parasite populations finally collapse and get extinct or rise jointly in size. The behavior of the allele frequency cycling is determined by the state of the fixed point (computed in SI 3 for *A* = 2). The cycling can be stable (regular cycles as fluctuating selection) or damp off to the fixed polymorphic state. The stability behavior of the hosts and the parasites, whose frequencies are not explicitly given by (1) and (2), cannot be explicitly determined by means of a Jacobian matrix and only be determined by numerically solving (1) and (2).

For the stability analysis, the initial conditions were chosen so that *N*^W^ is close to 10,000 and the infected alleles make up 20% of the healthy ones (*H*_1_ = *H*_2_ = 4150, *I*_11_ = *I*_12_ = *I*_21_ = *I*_22_ = 415). The birth rates b were fixed to one and *c_H_* = *c_P_* = 0.05 for the MA model and *c*_*H*_1__ = *c*_*P*_2__ = 0.05, *c*_*H*_2__ = *c*_*P*_1__ = 0 for the GFG model (see [20, 21] for comparable costs). The remaining parameters are given in the color-coded figures of SI 5 summarizing the results of the stability analysis, which become more diverse in behavior with a decreasing selection coefficient s, and an increasing difference between the birth rates *b* and the death rates *d*. We chose an appreciable number of initially infected alleles to omit cases were such genotypes could be instantaneously lost. Note that changing the birth rate to values other than one and employing various initial allele frequencies of healthy and infected alleles give results that correspond to the ones presented here (up to a rescaling of the underlying parameters).

The impact of the various parameters on the behavior of the dynamical system can be summarized as follows. An increasing difference between the birth and the death rates, *i.e.* between *b*(1 − *c_H_*) and *d*, results in 1) wider parameter ranges of the mortality *δ* and the disease transmission rate *β* for which cycles may occur, and 2) also increases the number of cycles over a given time interval. While the number of cycles generally increases with increasing values of *δ*, *β* affects the number of cycles most distinctly for *s* = 1, for which smaller values of *β* lead to a reduced number of cycles. The speed of cycling for MA and GFG models is equivalent, if costs are set to zero. Thus, it may be unrealistic to aim to infer the model itself (MA or GFG) based on polymorphism data. In the following, we focus on the GFG model and present according results for the MA model in the supplement.

As illustrated in Figure 2, besides the difference between *b*(1 − *c_H_*) and *d*, the selection coefficient *s* is the crucial parameter that determines the number of cycles per unit of time. A limit cycle is observed for *s* = 1, a case defined as “castrating parasite”. The cycling appears faster and with quicker damping off with smaller values of *s* [2]. Note that employing the same parameters to the MA model results in a loss of all parasite alleles for *s* = 1 and *s* = 0.9, whereas cycling towards the fixed point is obtained for *s* = 0.6 and *s* = 0.3. The MA model with a death rate *d* = 0.6 and otherwise equivalent parameters yields cycles for *s* = 1 and *s* = 0.9 as well (in contrast to the behavior when *d* = 0.9). The speed of cycling is therefore not only determined by the coevolutionary (*s, c_H_, c_P_, α*) and epidemiological parameters (*β, δ*), but also by the ecological characteristics of the species and the environment controlling the birth (*b*) and death (*d*) rates. An example of a MA model with a death rate *d* = 0.9 and slow cycles that will be studied alongside the GFG example of Figure 2 is given in SI 6.

**Figure 2.**
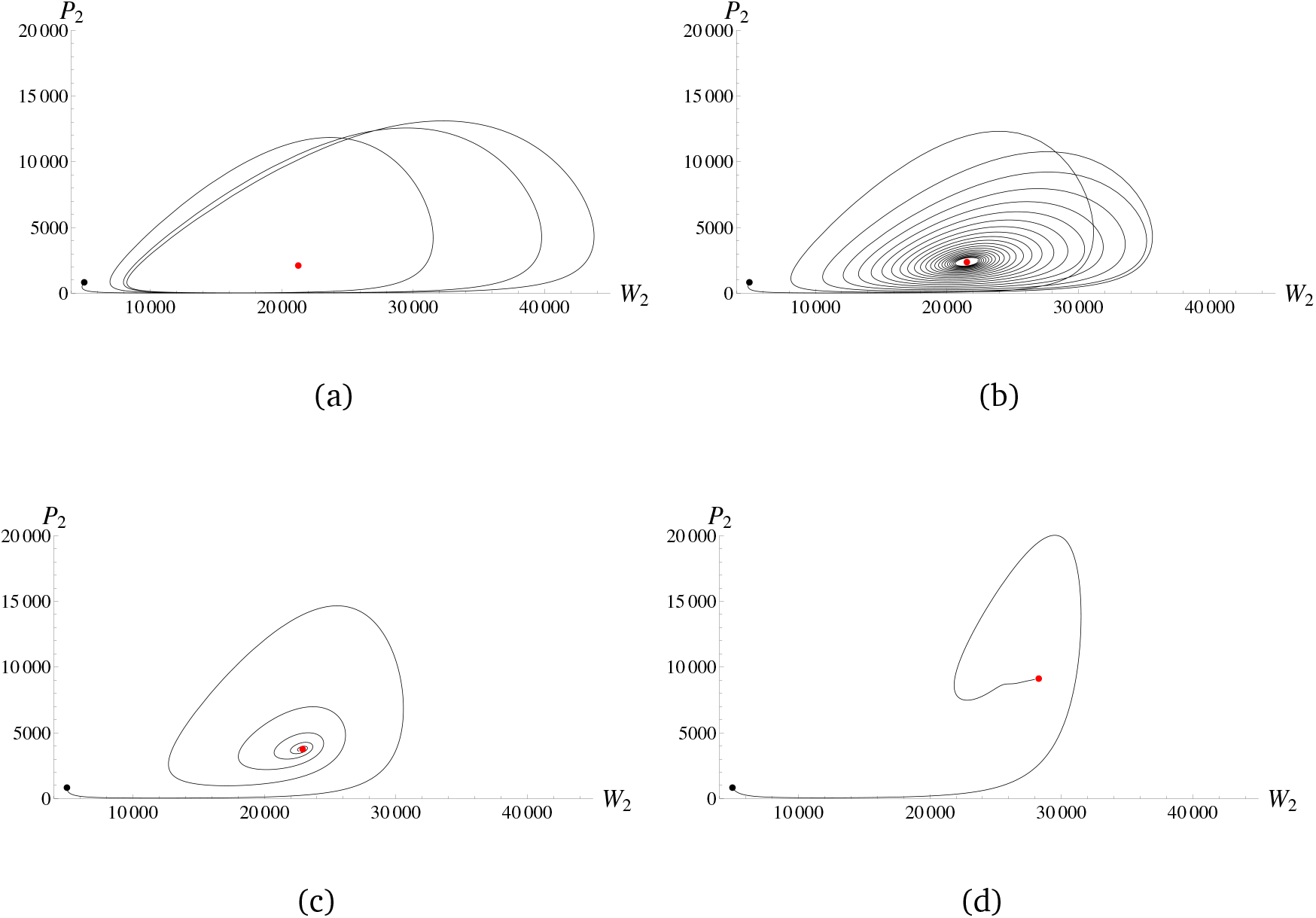
Parasite and host alleles of genotype two are plotted against each other for the GFG model over time by numerically solving (1) and (2) for the following parameter values (being equivalent for both genotypes): *b* = 1, *d* = 0.9, *δ* = 0.01, *β* = 0.00005, *c*_*H*_1__ = *c*_*P*_2__ = 0.05 and *c*_*H*_2__ = *c*_*P*_1__ = 0. The initial conditions are *H*_1_ = *H*_2_ = 4150 and *I*_11_ = *I*_12_ = *I*_21_ = *I*_22_ = 415. The selection coefficients *s* are given by (a) 1, (b) 0.9, (c) 0.6 and (d) 0.3. The parametric plots are shown for (a) 100, (b) 500, (c) 120 and (d) 96 time steps, which are the minimum amounts of time to complete one orbit of the final limit circle (a), come close to the fixed point (b), (c), or to reach the fixed point (d). The initial and fixed points are colored in black and red, respectively.

### Analyzing the polymorphism patterns of hosts and parasites

#### One parasite per host and equal generation times

We provide detailed results for a few examples of GFG and MA models to highlight the key features regarding changes in the host and parasite SFS. Mutation rate and genetic drift occur on different time scales for hosts and parasites. The population size changes in the parasite occur on a relatively slower time scale compared to the host (Figure 3). Moreover, the amplitude of population size fluctuations is much more pronounced than in the host, the number of infected alleles fluctuating between about twenty and 13.000 individuals. The relatively weak and fast fluctuations of the host cannot be observed in the SFS except for a slight increase in the number of singletons over time. In contrast, the strong and relatively slow changes in the parasite population size are clearly reflected by the same statistic. The number of singletons decreases first due to the initial decline in the population size before tending to increase over time. A similar result for the MA model is given in SI 7.

**Figure 3.**
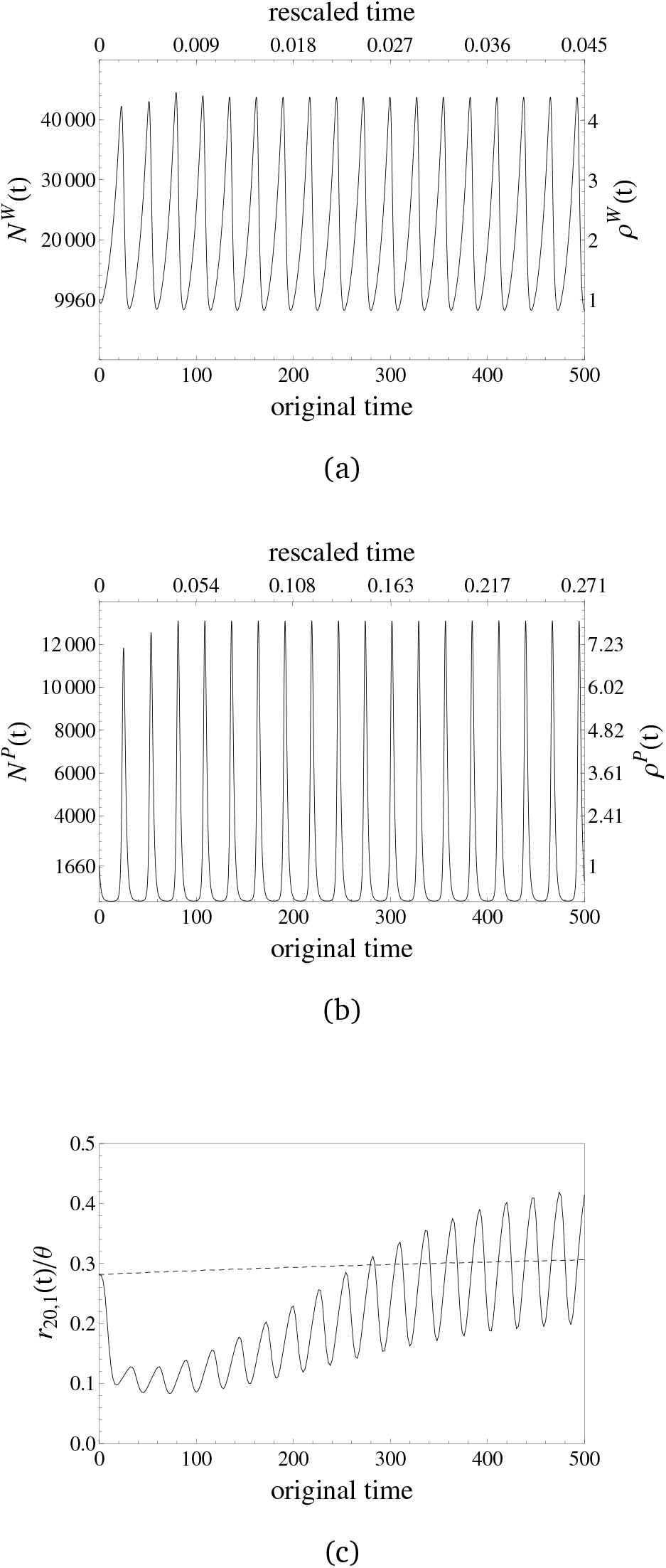
Population size changes in the host (a) and in the parasite (b) are generated for the GFG model via the parameters *b* = 1, *d* = 0.9, *δ* = 0.01, *β* = 0.00005, *c*_*H*_1__ = *c*_*P*_2__ = 0.05, *c*_*H*_2__ = *c*_*P*_1__ = 0, and *s* = 1. The initial conditions are *H*_1_ = *H*_2_ = 4150 and *I*_11_ = *I*_12_ = *I*_21_ = *I*_22_ = 415, so that the reference population sizes Nref of the host and the parasite are given by 9960 and 1660, respectively, before their interaction starts at time zero. In both cases, the lower x-axes show time at the original scale of the dynamical system, whereas time is scaled by the respective values of *N*_ref_ · 1/*d* for the upper x-axes. The left y-axes denote the absolute values of the changing population size and the right y-axes denote the relative values as *ρ*(*t*) = *N*(*t*)/Nref. We evaluated the SFS for every second generation (on the original scale) in both cases and plot (c) the relative number of singletons *r*_20,1_(*t*) against time for the host (dashed) and the parasite (solid).

We also observed that signatures of coevolutionary cycles in the host and parasite SFS depend mostly on the speed of their fluctuations in terms of population size scaled generation times, whereas the magnitude of the population size changes has a small influence on the SFS (rather apparent in Figure 1 than in Figure 3). We also evaluated one of the slowest cycling examples for a death rate of *d* = 0.3 shown in the first panel in Figure SI 5.2.2 to illustrate that despite the difference in time scale in hosts and parasite cycles are also barely detectable in the parasite SFS (SI 8).

To detect changes in the SFS in host and parasite, it is also important to assess if enough genetic diversity can be observed over time. We find that the absolute number of polymorphisms strongly decreases over the considered time interval. For example in the GFG model (Figure 3) the number of segregating sites in the parasite decreases to about five percent and for the MA model (SI 7) even down to about one percent of their initial values. We also contrast two scenarios: 1) an initial total host population size of 100,000 and 2% of initial disease prevalence, and 2) an initial population size of 10,000 with 50% initial prevalence (for GFG and MA models in SI 9). We observe that the initial prevalence defining the parasite population size plays a crucial role. Scenario two shows stronger fluctuations in the relative number of parasite singletons and a more drastic decrease in the absolute number of segregating sites over time than under scenario one. This shows that larger fluctuations in the relative SFS over time, which are in principle detectable in time sample polymorphism data, go along with a stronger decrease in the total amount of observed polymorphisms (due to more drastic population bottlenecks). As a guideline, we provide an estimate of the possibility to detect changes in the SFS based on a sufficient number of segregating sites (the numerical minima and maxima in Figure SI 9.1 yield Table 2). Cycles are more likely observed in parasites with large genome mutation rates such as fungi or protozoans in contrast to bacteria (Table 2).

**Table 2.** [minimum, maximum] of the number of segregating sites in full genome sequences of parasites following the GFG model and depending on the initial prevalence and population size. The minima are determined by the results in SI 9 and the maxima are the initial numbers of segregating sites. The per genome mutation rate, *μ*, is an approximation based on genome length and per site mutation rate from different typical estimates [22].

| Initial prevalence,<br>$N_{\text{ref}}$ parasites | Fungi/Trypanosoma/Nematodes<br>$\mu = 1$ | Viruses<br>$\mu = 0.1$ | Bacteria<br>$\mu = 0.001$ |
| --- | --- | --- | --- |
| 2%, 2000 | [8232, 14191] | [823, 1419] | [8; 14] |
| 20%, 1660 | [560; 11779] | [56; 1178] | [1; 12] |
| 50%, 3332 | [48; 23642] | [5; 2364] | [0; 24] |

When comparing our computationally advantageous approach based on the Wright-Fisher model with our stochastic simulations, we find that polymorphism signatures as exemplarily measured by Π_20_(*t*) agree in general for host and parasite over time (Figure 4). The parasite sample shows less polymorphism in the simulations with overlap while for the host this difference is negligible. The net effect of generation overlap under strong population bottlenecks with more dominant decline than expansion phases is an even stronger decrease of the effective population size and thus the amount of polymorphism, as seen in the parasite. This is due to less newborns contributing fewer novel mutations in the model with overlap and the different sampling schemes of newborn and overlapping individuals. Whenever population size decreases, new offspring individuals are present in smaller proportions than overlapping ones, whereas the reverse occurs when population size increases. The fraction of new offspring follows a sampling with replacement as for the Wright-Fisher model, while the overlap fraction is drawn without replacement. The difference of these two sampling schemes becomes apparent during phases of small population sizes. During decline (expansion) phases, the increased fraction of overlapping (newborn) individuals leads to stronger (lesser) deviations from the Wright-Fisher expectations. Consequently, as the parasite population is experiencing a drastic population decrease over time and several cycles, the amount of diversity differs between both approaches in contrast to the host population experiencing a slight increase in size over time.

**Figure 4.**
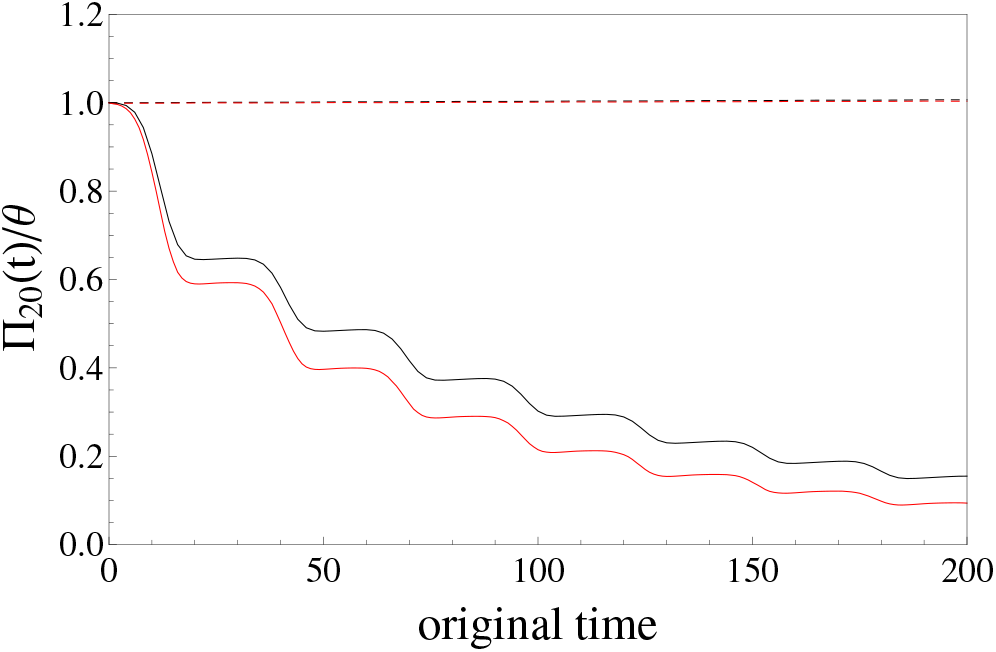
Population size changes in host (dashed) and parasite (solid) are generated for the GFG model with the same parameter set as in Figure 3. The average number of pairwise differences scaled by *θ*, Π_20_(*t*)/*θ*, is plotted against time for our analytical framework based on the diffusion approximation of the Wright-Fisher model (black) and for our simulation approach (red) with a fractional overlap of 1 − *d* (*d* = 0.9) between successive time steps. For the simulations, computational time steps are set to 0.001, a genomewide mutation rate of *μ* = 1 is applied, and each value is obtained as an average over 10 repetitions.

#### Multiple parasites per host and polycyclic diseases

We extend our predictions for two classic deviations from our model. First, multiple or even a large number of parasites, as denoted by *F*, often infect a single host. Considering this effect on the SFS leads to an increased scaled mutation rate *F θ* thereby increasing the number of segregating sites that can be detected by full-genome sequencing. Concurrently, relative time is sped up by *F* due to the larger initial (reference) population size of the parasite. Therefore, two opposite effects are expected when increasing the number of parasites per host, an increase in the amount of polymorphism comes at the cost of a reduced amount of detectable cycles. Second, host and parasite generation times may differ from one another with parasites often exhibiting smaller generation times especially for virus, bacteria or fungi (polycyclic diseases). We define *E* parasite generations per host generation so that the relative time for the parasite is slowed down by *E*. This rescaling is expected to enhance the detectability of coevolutionary cycles using the parasite SFS. We investigate the joint impact of multiple parasites per host and polycyclic diseases in SI 10.

We compute the SFS over time for the GFG above (Figure 3) with nine different combinations of values for *E* and *F* (see SI 10). To evaluate fluctuations in the polymorphism pattern over time, we compute the relative rate of change of Π_20_(*t*) (SI 10) at various equidistantly distributed sampling points over time. These points are chosen to be fixed in time and independent of parasite generation time, so that the various cases are equivalently clocked on the original time scale of the dynamical system. As illustrated in SI 10 more sampling points are needed to capture the cycling for multiple parasites per host (F), whereas less samples are needed for polycyclic diseases. We therefore show here that the number of time samples necessary to recover the number of cycles can be determined knowing the biology of host and parasite.

## Discussion

We develop here a model to analyze the evolution of neutral genome-wide polymorphism of coevolving host and parasite populations. Variations in polymorphism reflect the co-demographic history driven by eco-evolutionary feedbacks between 1) the frequency changes in host resistance and parasite infectivity over time due to frequency-dependent selection, and 2) the ecological changes in host and parasite population sizes. While antagonistic or synergistic coevolution is a process driven by natural selection, our approach is the first to provide a description of the consequences of coevolutionary dynamics on neutral genome-wide polymorphism.

We demonstrate that using time sampling data, *i.e.* population samples of hosts and parasites at different time points, it is possible to track the existence and speed of eco-evolutionary cycles using polymorphism data. Our main result states that cycles of coevolution are detectable in the parasite and barely in the host population. This is due to a fundamental difference in the time scale of neutral evolution between interacting species and the strength of their size fluctuations over time both crucially depending on the initial population size at the onset of epidemics (*i.e.* the start of the coevolutionary history). The time scale of neutral evolution is also determined by the parasite generation time and number of parasites per infected host. We study here one coevolutionary run starting by the introduction of a parasite population into a larger host population and generating a dynamics over several hundreds of generations. If new infectivity or resistance alleles appear by mutation, the epidemics and the cycling behavior are affected, and our model should be reset to evaluate a new run. The time scale that we investigate is therefore intermediate between the classic expectations of long coevolution and its signatures at interacting loci [6, 23] and the short-term epidemiological dynamics (with susceptible hosts and one parasite type) [24].

Polymorphism data can be used to detect coevolutionary cycles, if such cycles run sufficiently long at adequately low speed. We indeed predict that long term occurrence of cycles should be searched for in parasites that strongly decrease the host fitness due to high disease severity *s* (parasite effect on fecundity). For low to moderate disease transmission and smaller values of *s* the internal polymorphic equilibrium point is a stable attractor, meaning that cycles damp off quickly towards a polymorphic equilibrium at which population size and allele frequencies are fixed (Figure 2). For high values of the disease transmission rate *β* and parasite virulence *δ* (effect of parasite on mortality), the internal polymorphic equilibrium point is an unstable point [2], and a monomorphic equilibrium is reached with a fixed population size [10, 12]. Cycles should be slow enough to be observable in polymorphism data, and our results challenge the classic assumption that coevolutionary cycles are too fast for being observed in the SFS. Interestingly, the speed of cycling depends mainly on two ecological parameters, *i.e.* the birth rate *b* and the death rate *d* of the host, and to a lesser extent on the coevolutionary parameters (*s*, costs of resistance and virulence).

Eco-evolutionary cycles occur and are observable in polymorphism data, when the effect of the parasite on host fecundity (*s*) is sufficiently strong. We predict that our results are applicable to many host-parasite systems with castrating parasites, whose transmission is associated with host death (algae-rotifer [25], bacteria–phage [26] and *Daphnia magna*–bacterium [27]). For plant pathogens, cycles may be less observable because the disease severity can range from low to very high (or even castrating, [28]), but often depends on abiotic factors [29]. Most epidemiological studies have focused on the evolution of virulence (effect of parasite on host mortality) and disease transmission within the short duration of an epidemics. However, to use polymorphism data for the study of eco-evolutionary dynamics, parasite virulence is not an essential parameter to be measured or estimated, as it is more useful to quantify the difference between the host’s birth and natural death rates. Another practical reason for favoring hosts with high death rates is that our SFS computations are based on the Wright-Fisher model assuming non-overlapping generations [15]. Note that in epidemiological models [2, 10, 12] overlapping generations in the host are a necessary assumption, since a disease can only be transmitted among living hosts. We wrote a code in C (available from the authors upon request) that explicitly accounts for less newborns contributing fewer mutations while generations overlap (and compared to the Wright-Fisher model) to evaluate the SFS. The simulations show that our Wright-Fisher approximation is robust with respect to overlapping generations for hosts with high death rates.

We only consider neutral sites because arbitrary demographic changes can be used in the analytic solution for the SFS as needed to cope with the complex demographies arising from our dynamical system. The frequency spectrum for sites under selection can only be computed for piecewise changes in the population size [30]. This approach is not readily applicable for such complex demographies because their discretization would be computationally cumbersome.

Time sampling is crucial for capturing cycles (see SI 10) that can be observed in the polymorphism data for several hundreds of parasite generations, if the genome mutation rate and the effective population size are sufficiently large (see Table 2) and if SNPs are sampled at appropriate time points (see SI 10). To test our predictions, time samples can be readily obtained in experimental coevolution set-ups [25, 26], whereas this may be more complex for natural populations. Nevertheless, samples from the past can be obtained for crustaceans (*Daphnia*, [27]) from dormant stages deposited in sediments, and for plant species from seeds in the soil (possibly using ancient DNA recovery techniques).

We predict that parasites undergoing several generations per host generation but producing small amount of pathogen propagules per host should be the species exhibiting most clearly the signature of co-demographic dynamics in polymorphism data. Our results pave the way to use time sample genomic data of hosts and parasites from wild or experimental populations, to analyze, infer and take into account the co-demographic history of the antagonistic species and scan for genes under coevolution with greater accuracy.

## Supplementary Information

### SI 1. The host effective population size over time

From (1) and (2) we immediately obtain

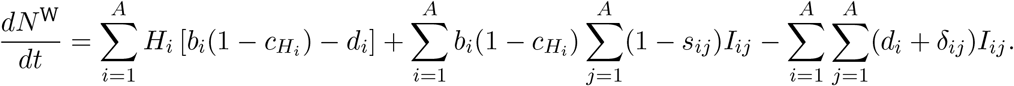

Assuming that *s_ij_ = s_i_, δ_ij_ = δ_i_*(*i.e.* both parameters being independent of the parasite genotype) and setting *dN*^W^/*dt* to zero, we have

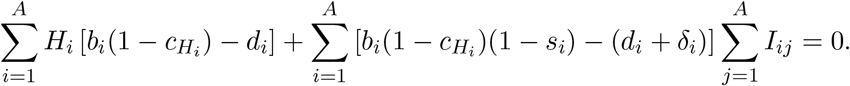

It particularly follows that *s_i_ = δ_i_* = 0 requires *d_i_ = b_i_*(1 − *c_H_i__*) to have a constant population size in the host for arbitrary choices of *H_i_* and *I_ij_*.

### SI 2. Basic reproduction ratios

From (2), we obtain

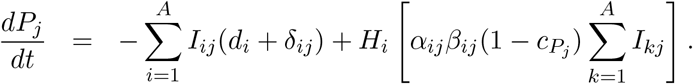

Assuming that *d_i_ = d* and *δ_ij_ = δ_j_*(*i.e.* both parameters being independent of the host genotype), the former equation simplifies to

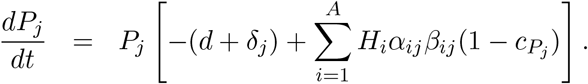

The reproduction ratios of the parasite genotypes are given by

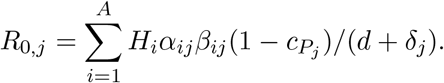

If all *R*_0,*j*_ < 1, the parasite genotypes are eliminated since they kill more hosts than they infect new healthy ones. When one of these ratios is greater than one, the corresponding parasite genotype is maintained in the population.

### SI 3. Fixed points of the dynamical system

We calculated the fixed points by setting (1) and (2) to zero. Note that the solutions were first obtained in terms of the numbers of healthy and infected hosts before being added up to obtain the fixed points of hosts and parasites assuming one parasite per host. The results for the two-allele case are summarized below. For the MA and iGFG model the results for more than two alleles correspond to those of two alleles, whereas results for the iMA and the GFG model can only be obtained for special cases when *A* = 3, but the solutions are not enlightening.

Define 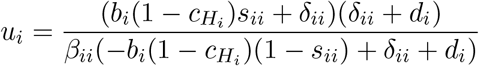 and 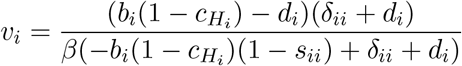.

For the **MA model** the fixed point, where all alleles may have nonzero frequencies, is given by (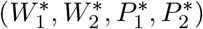), where for *i* = 1,2,

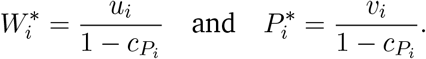

Besides the trivial solution of all alleles having frequency zero, the remaining solutions are given by (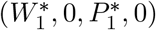) and (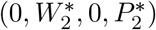).

For the **iMA model** the fixed point, where all alleles may have nonzero frequencies, is given by (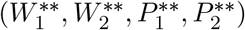), where for *i* = 1,2,

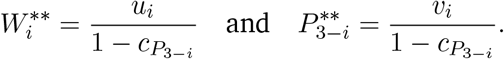

Besides all alleles having frequency zero, the remaining solutions are given by (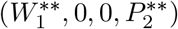) and (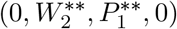).

For the **GFG model** the fixed point, where all alleles may have nonzero frequencies, can be evaluated but is of complicated form. So we only note the results for *β_ij_ = β_i_, δ_ij_ = δ_i_* and *s_ij_ = s_i_*(*i.e.* all of these parameters are independent of the parasite genotype). The fixed point with nonzero entries is given by (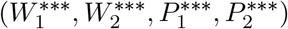), where

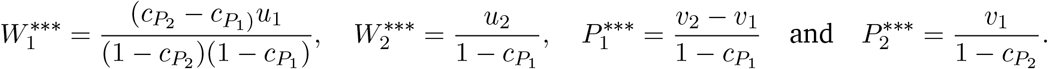

Besides all alleles having frequency zero, further solutions are given by (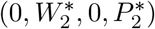), (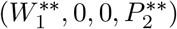) and (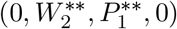).

For *c*_*H*_1__ = *c*_*H*_2__ = *c*_*P*_1__ = *c*_*P*_2__ = 0, we also obtain

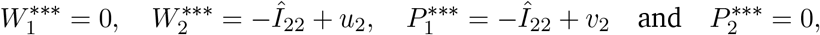

where 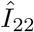 denotes the equilibrium solution of the infected host genotype. With the additional assumption of equivalent rates and costs among both host and parasite genotypes, so that particularly *u*_1_ = *u*_2_ = *u* and *υ*_1_ = *υ*_2_ = *υ*, we further have

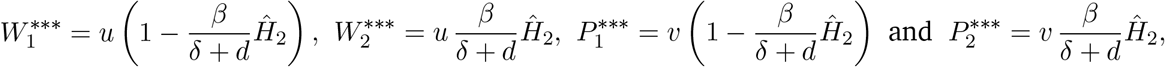

where 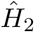 denotes the equilibrium solution of the second healthy host genotype.

For the **iGFG** model, the nontrivial equilibrium solution is equivalent to the MA model with 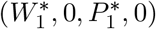.

### SI 4. Summary of analytical results for site frequency spectra and related statistics

Assume a model, where a haploid population is evolving according to Wright-Fisher dynamics forwards in time and being of size *N*_ref_ at (and before) time zero. A sample of *n* sequences is taken. Neutral mutations occur at unlinked and previously monomorphic sites at rate *θ* = 2 *N*_ref_ *μ, μ* being the mutation rate per genome per generation. Scale time in units of *N*_ref_ generations and let *N*_ref_ → ∞ to reach the diffusion limit. Thereby, the relative population size *N*(*t*)/*N*_ref_ converges to the strictly positive and piecewise continuous scaling function *ρ*(*t*). The site frequency spectrum is the distribution of the number of times a mutant allele is observed in the sample among the polymorphic loci.

The absolute site-frequencies over time *f_n,i_*(*t*), 1 ≤ *i* ≤ *n* − 1, with a mutation-drift equilibrium at time zero can be simply obtained from Equation 33 in [15] as

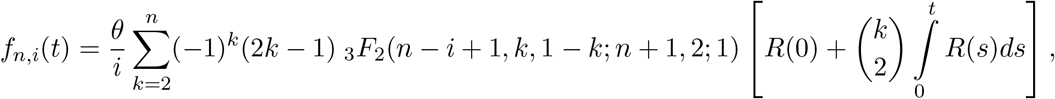

where 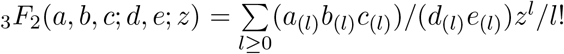 for *l* ≥ 1, is a *generalized hypergeometric function*, and 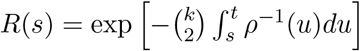.

The relative site frequencies over time *r_n,i_*(*t*) are obtained as 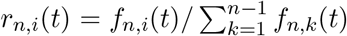, where the denominator gives the absolute number of segregating sites, *S_n_*(*t*). For the average number of pairwise differences, we have 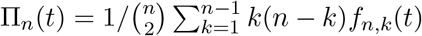.

These implementations can be computationally demanding since the inverse of the relative population size function *ρ*(*t*) has to be integrated numerically with high precision before being applied to an exponential function that has to be integrated numerically again from the initial time zero up to the time points at which the SFS are evaluated. These exponentials also cover binomial coefficients that depend on sample size and the numerical integration has to be performed for every single one of those. Therefore, an increasing amount of integration steps and computation time is required with increasing sample size, so that we only consider a rather small sample size of twenty individuals for which the SFS of a single time point can be rather quickly obtained and even for later time points. This is particularly desirable, since we are interested in taking samples recurrently over the course of time.

### SI 5. Stability behavior of the fixed points for two alleles

#### SI 5.1. The matching-allele model

**Table SI 5.1.1.**
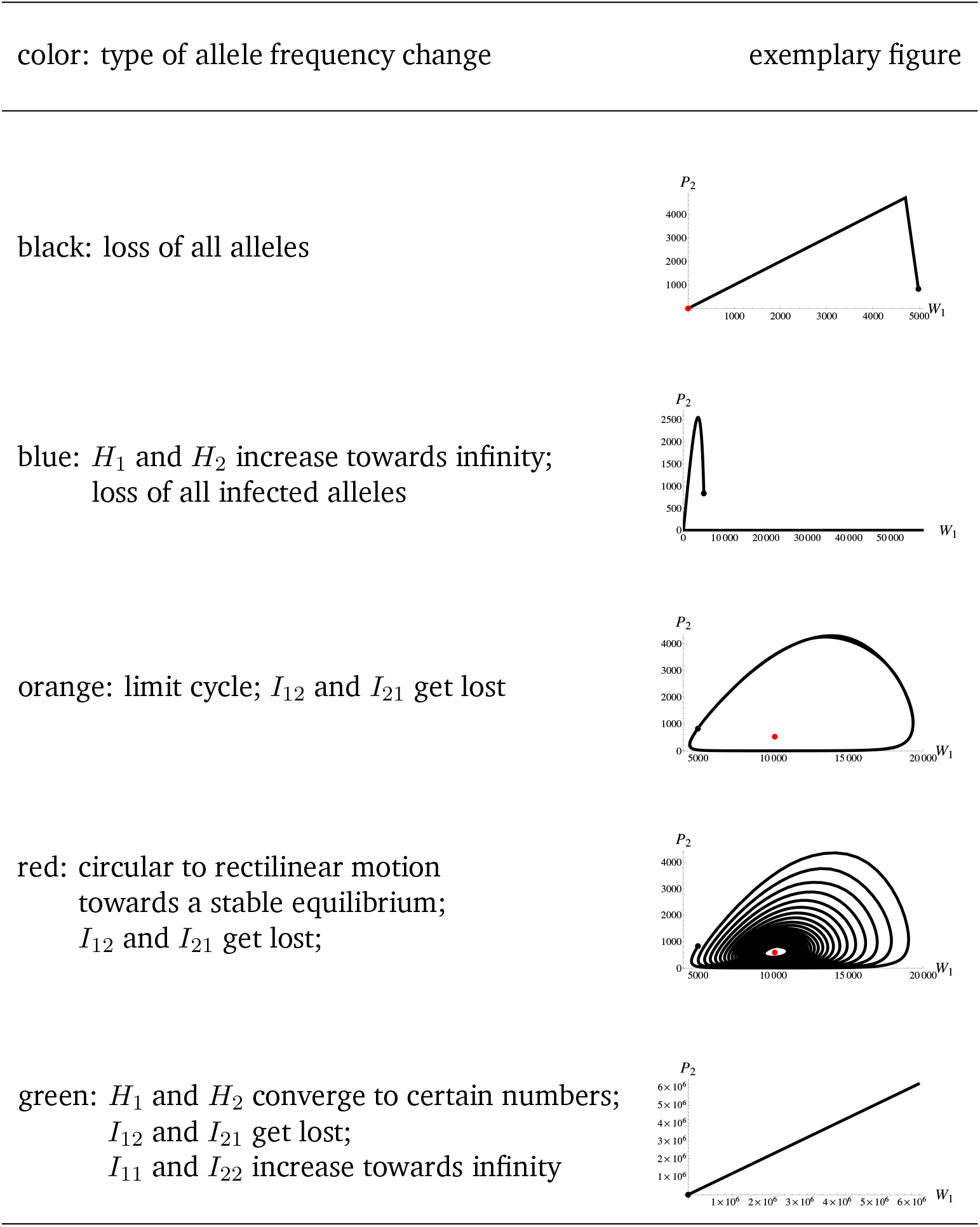
Legend of coloring for possible allele frequency changes. In the left column the possible fates of (healthy and infected) genotypes are summarized and assigned to colors. In the right column corresponding exemplary parametric plots of host and parasite allele frequencies are shown using Equations (1) and (2). The black dot depicts the initial allele frequency, whereas the red dot shows the zero point in the first and the fixed point in the third and in the fourth subfigure.

| color: type of allele frequency change | exemplary figure |
| --- | --- |
| black: loss of all alleles |  |
| blue: $H_1$ and $H_2$ increase towards infinity;<br>loss of all infected alleles | |
| orange: limit cycle; $I_{12}$ and $I_{21}$ get lost | |
| red: circular to rectilinear motion<br>towards a stable equilibrium;<br>$I_{12}$ and $I_{21}$ get lost; | |
| green: $H_1$ and $H_2$ converge to certain numbers;<br>$I_{12}$ and $I_{21}$ get lost;<br>$I_{11}$ and $I_{22}$ increase towards infinity | |

**Figure SI 5.1.1.**
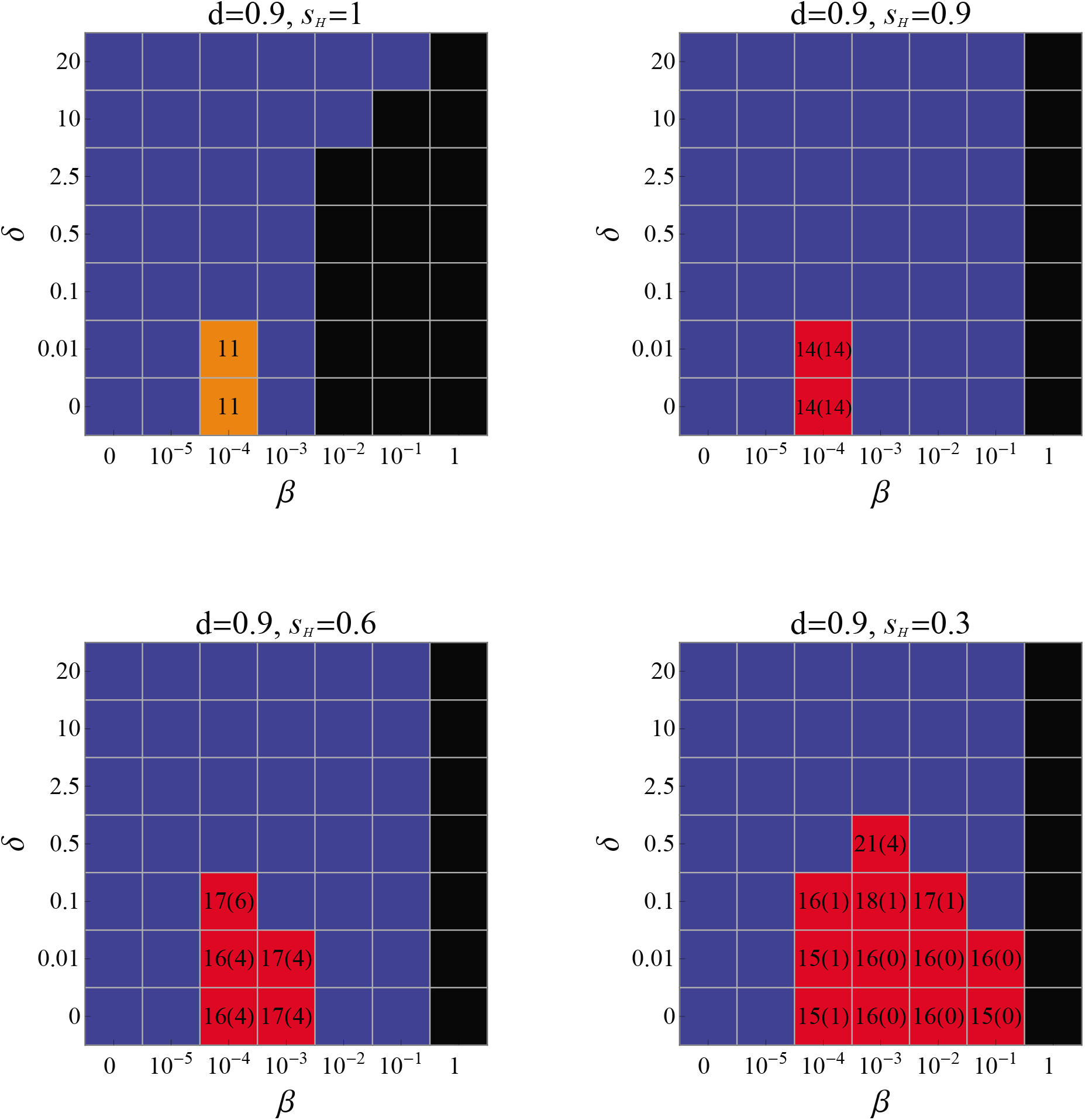

**Figure SI 5.1.2.**
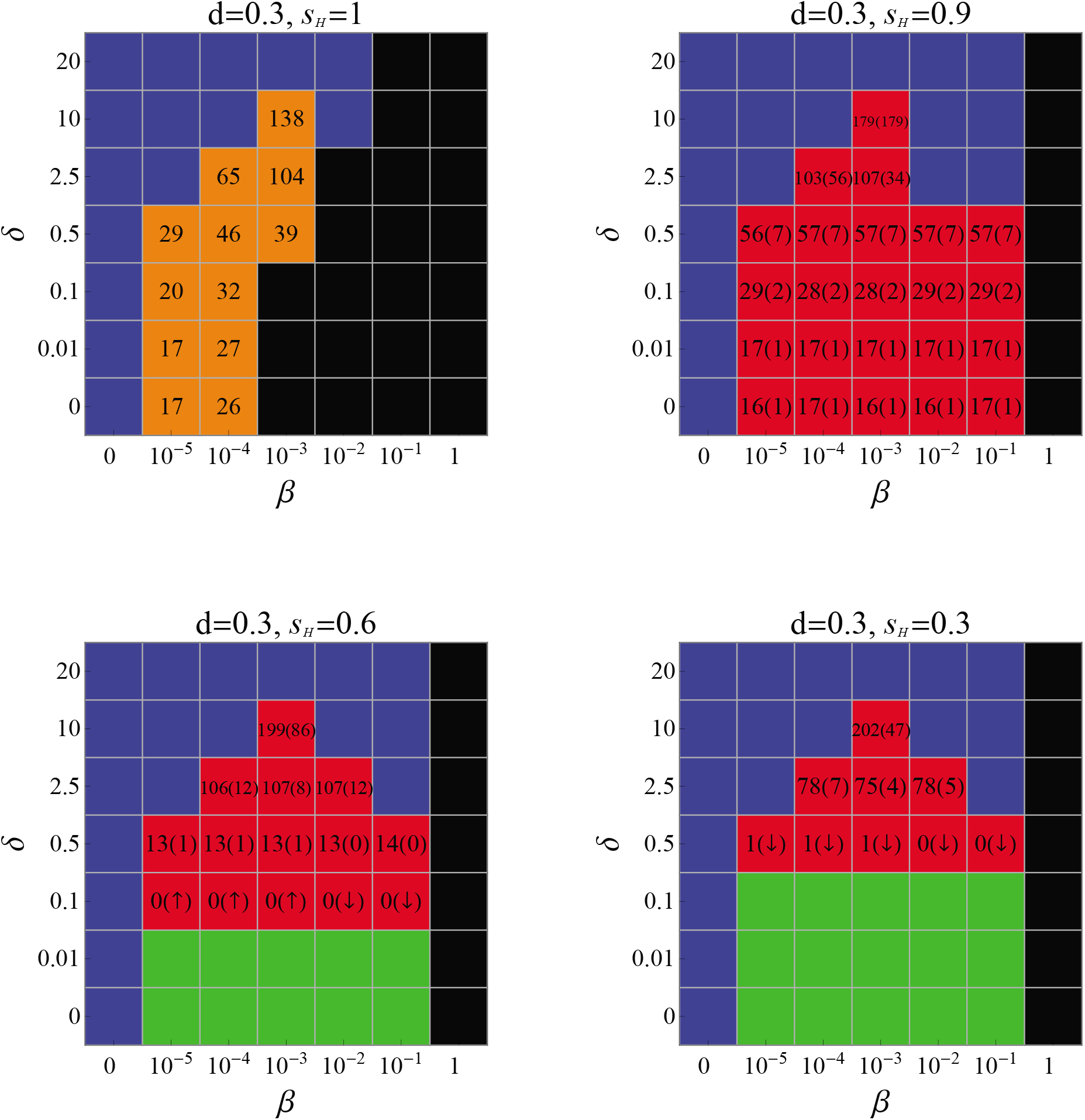

#### Description of the results

For each panel of Figures SI 5.1.1 and SI 5.1.2 the system of differential equations (1) and (2) were numerically evaluated for *b* = 1, *c_P_ = c_H_* = 0.05 and the specified parameters over 500 time steps. The various possible coevolutionary scenarios of hosts and parasites are color-coded and summarized in detail in Table SI 5.1.1. The squares featuring numbers represent the parameter combinations for which cycling around or immediate attainment of the fixed point of *N*^W^ is obtained. For *s* = 1, the amplitude of the cycles around the fixed point of *N*^W^ remains constant over time. For other values of *s* we distinguish two cases: We first evaluated the total amount of cycles around the fixed point of *N*^W^ based on a numerical accuracy of 20 digits. Since the cycling dumps off in these cases, we also note in brackets the number of cycles until the amplitude is enclosed by the interval [0.95 *N*^W^, 1.05 *N*^W^]. The upwards and downwards arrows in some of the brackets illustrate expansions and declines to the fixed point, respectively.

#### SI 5.2. The gene-for-gene model

**Table SI 5.2.1.**
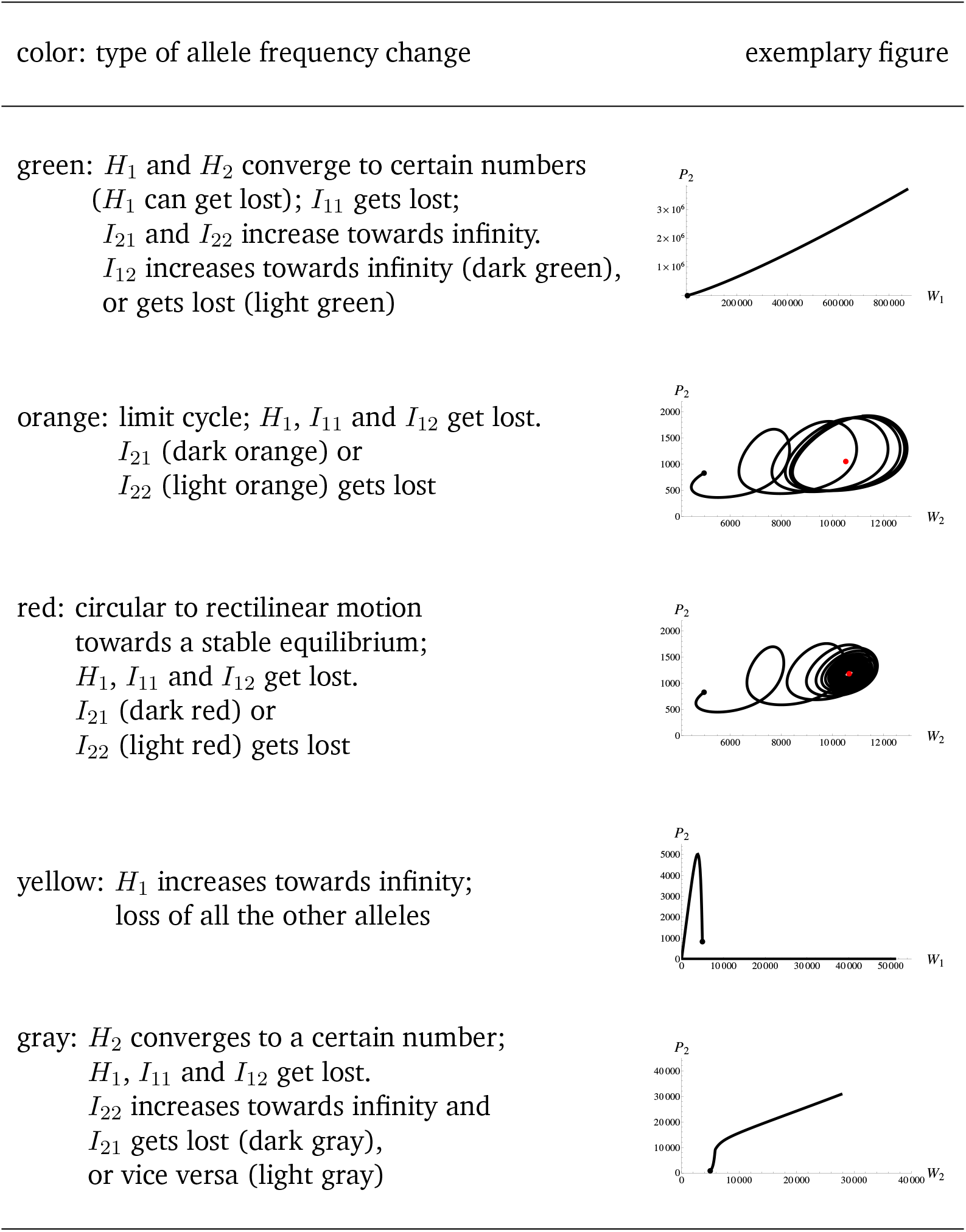
Legend of coloring for possible allele frequency changes. In the left column the possible fates of (healthy and infected) genotypes are summarized in addition or slightly modified to the scenarios already presented in Table SI 5.1.1 by means of the MA model. In the right column exemplary parametric plots of host and parasite allele frequencies are shown (in the first three panels for the dark color). The black dot depicts again the initial allele frequency.

| color: type of allele frequency change | exemplary figure |
| --- | --- |
| green: $H_1$ and $H_2$ converge to certain numbers ($H_1$ can get lost); $I_{11}$ gets lost; $I_{21}$ and $I_{22}$ increase towards infinity. $I_{12}$ increases towards infinity (dark green), or gets lost (light green) |  |
| orange: limit cycle; $H_1$, $I_{11}$ and $I_{12}$ get lost. $I_{21}$ (dark orange) or $I_{22}$ (light orange) gets lost |  |
| red: circular to rectilinear motion towards a stable equilibrium; $H_1$, $I_{11}$ and $I_{12}$ get lost. $I_{21}$ (dark red) or $I_{22}$ (light red) gets lost |  |
| yellow: $H_1$ increases towards infinity; loss of all the other alleles |  |
| gray: $H_2$ converges to a certain number; $H_1$, $I_{11}$ and $I_{12}$ get lost. $I_{22}$ increases towards infinity and $I_{21}$ gets lost (dark gray), or vice versa (light gray) |  |

**Figure SI 5.2.1.**
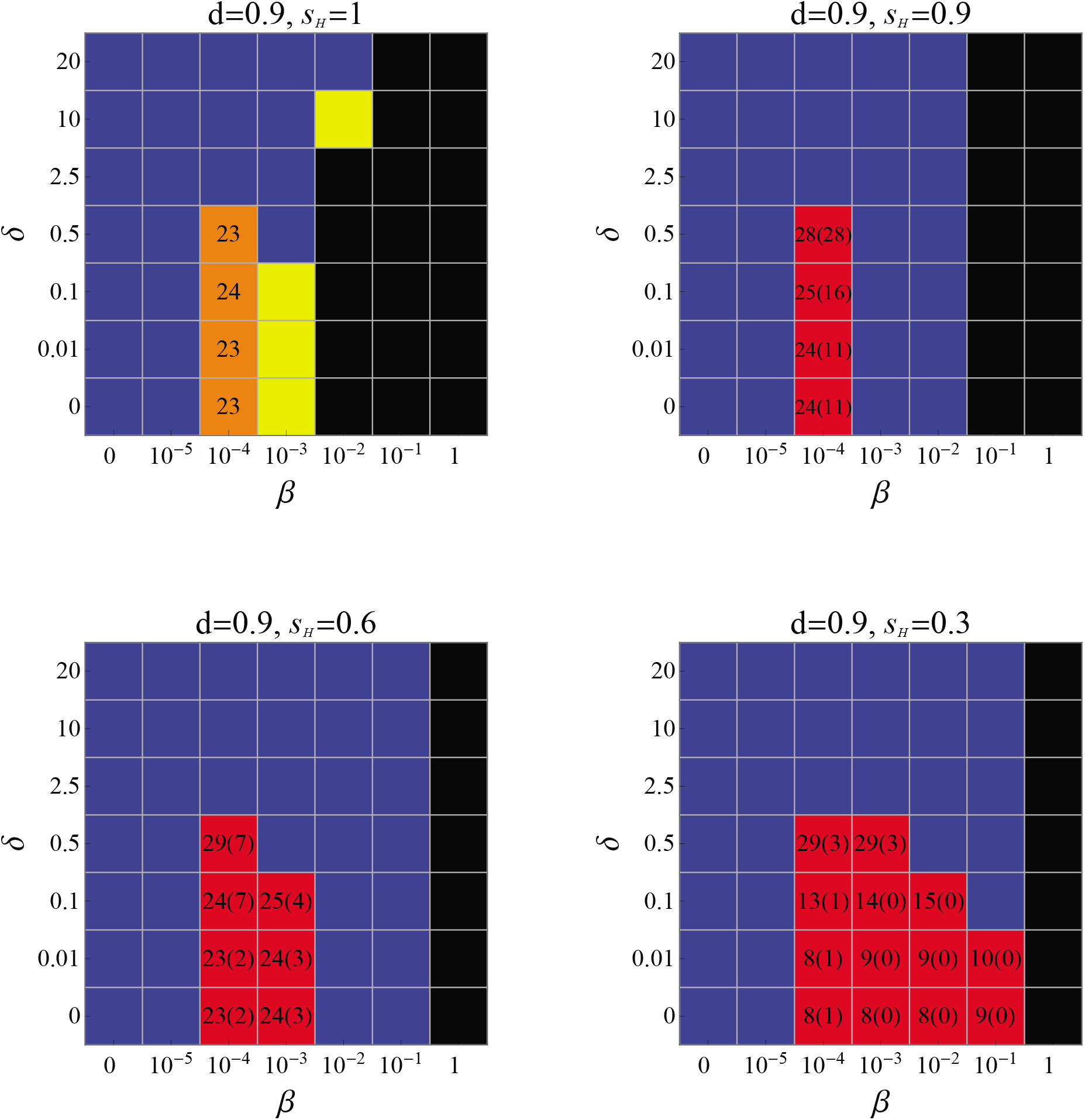

**Figure SI 5.2.2.**
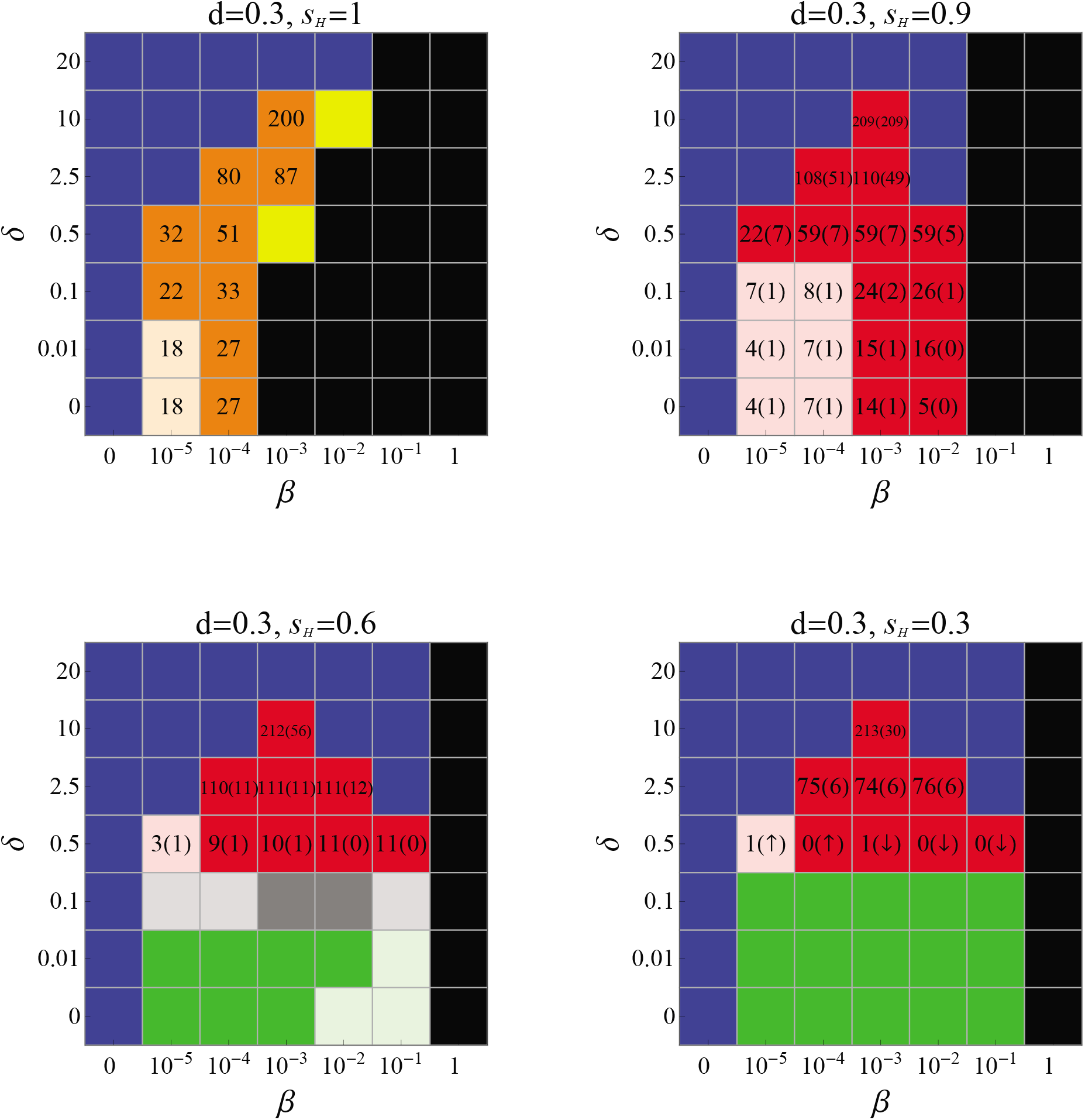

##### Description of the results

For each panel of Figures SI 5.2.1 and SI 5.2.2 the system of differential equations (1) and (2) were numerically evaluated for *b* = 1, *c*_*H*_1__ = *c*_*P*_2__ = 0.05, *c*_*H*_2__ = *c*_*P*_1__ = 0 and the specified parameters over 500 time steps. The various possible coevolutionary scenarios of hosts and parasites are color-coded and summarized in detail in Tables SI 5.1.1 and SI 5.2.1; the latter table summarizes cases either slightly different for or even exclusive to the GFG model due to the additional possibility of infection. The squares featuring numbers represent the parameter combinations for which cycling around or immediate attainment of the fixed point of *N*^W^ is obtained. For *s* = 1, the amplitude of the cycles around the fixed point of *N*^W^ remains constant over time. For other values of *s* we distinguish two cases: We first evaluated the total amount of cycles around the fixed point of *N*^W^ based on a numerical accuracy of 20 digits. Since the cycling dumps off in these cases, we also note in brackets the number of cycles until the amplitude is enclosed by the interval [0.95 *N*^W^, 1.05 *N*^W^].

### SI 6. Parametric plots for a matching-allele example

**Figure SI 6.1.**
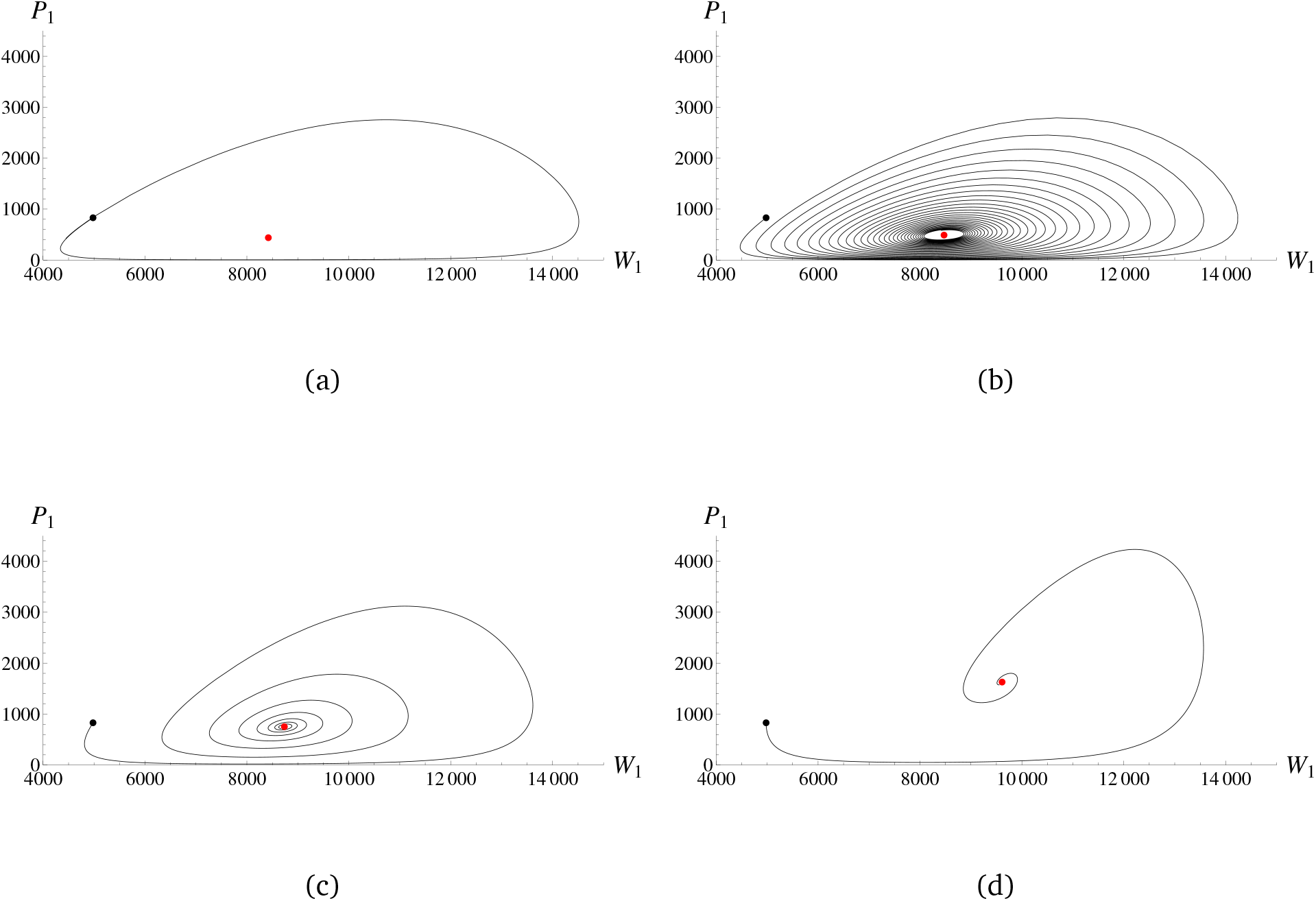
Parasite and host alleles of genotype one are plotted against each other for the MA model over time by numerically solving (1) and (2) for the following parameters (being equivalent for both genotypes): *b* = 1, *d* = 0.9, *δ* = 0.01, *β* = 0.00012 and *c_P_* = *c_H_* = 0.05. The initial conditions are *H*_1_ = *H*_2_ = 4150 and *I*_11_ = *I*_12_ = *I*_21_ = *I*_22_ = 415. The selection coefficients *s* are given by (a) 1, (b) 0.9, (c) 0.6 and (d) 0.3. The parametric plots are shown for (a) 40, (b) 900, (c) 225 and (d) 76 time steps, which are the minimum amounts of time to complete one circle (a), come close to the fixed point (b), or to reach the fixed point (c), (d). The initial and fixed points are respectively colored in black and red.

### SI 7. Time scaling and the distribution of singletons over time for a matching-allele example

**Figure SI 7.1.**
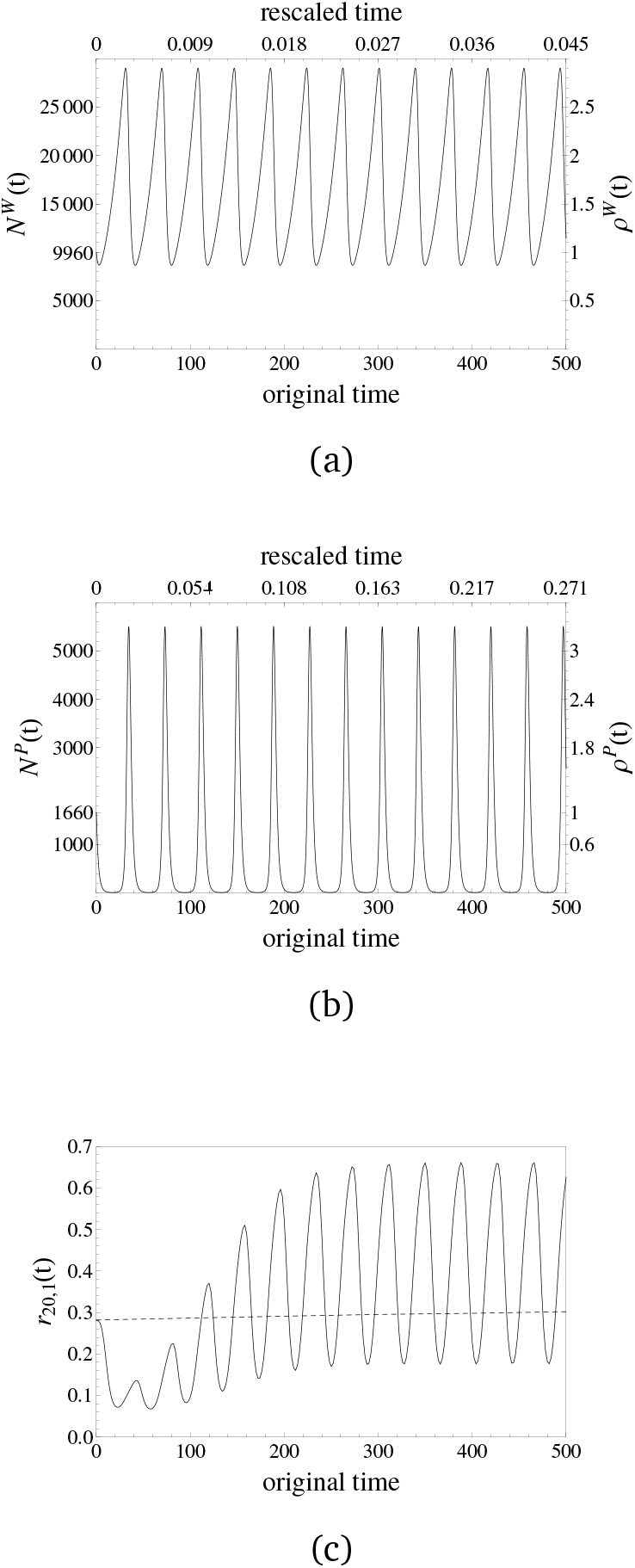
Population size changes in the host (a) and in the parasite (b) are generated for the MA model via the parameters *b* = 1, *d* = 0.9, *δ* = 0.01, *β* = 0.00012, *c_P_* = *c_H_* = 0.05, and *s* = 1. The initial conditions are *H*_1_ = *H*_2_ = 4150 and *I*_11_ = *I*_12_ = *I*_21_ = *I*_22_ = 415, so that the reference population sizes Nref of the host and the parasite are respectively given by 9960 and 1660 before their interaction starts at time zero. In both cases, the lower x-axes show time at the original scale of the dynamical system, whereas time is scaled by the respective values of Nref · 1/*d* for the upper x-axes. The left y-axes denote the absolute values of the changing population size and the right y-axes denote the relative values as *ρ*(*t*) = *N*(*t*)/*N*_ref_. The number of infected alleles fluctuate between about ten and 5.500 individuals. We evaluated allelic spectra for every second generation (on the original scale) in both cases and (c) plot the relative number of singletons *r*_20,1_(*t*) against time for the host (dashed) and the parasite (solid).

### SI 8. Time scaling and the distribution of singletons over time for a gene-for-gene example

**Figure SI 8.1.**
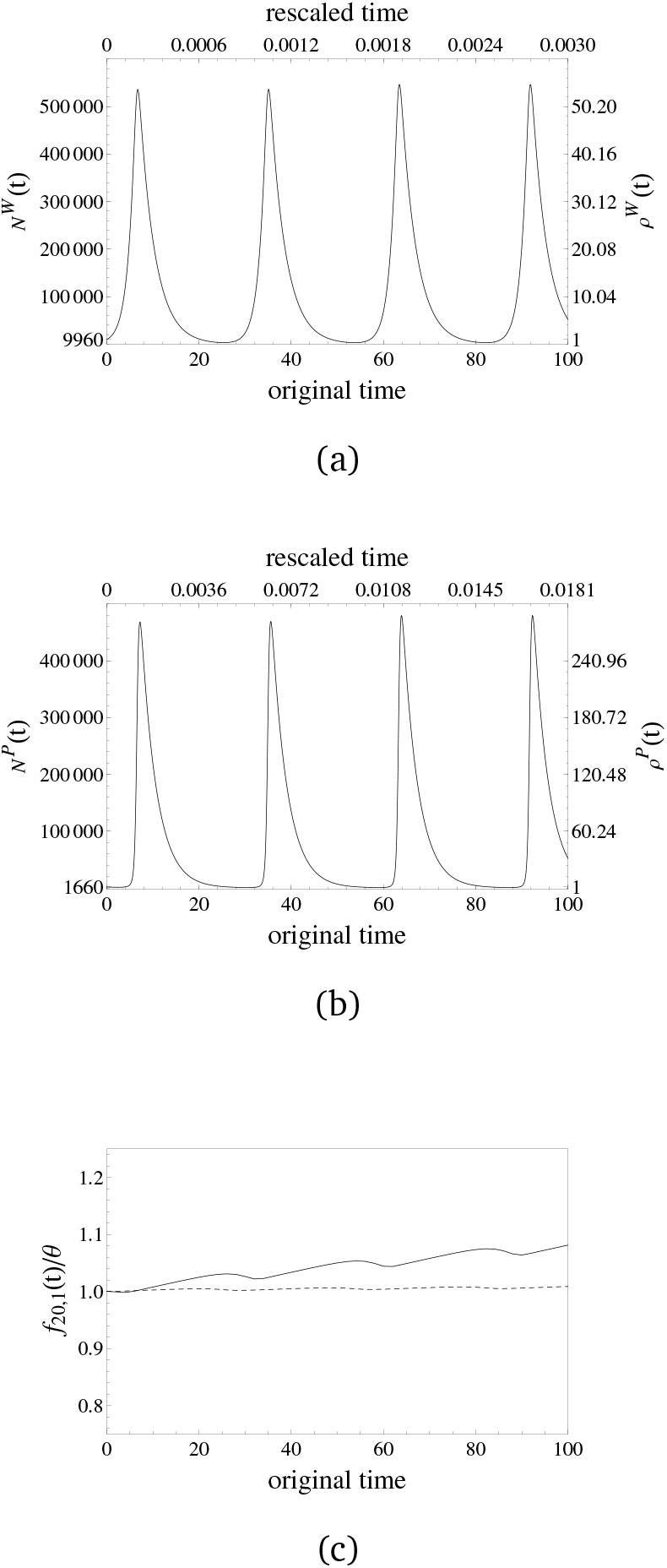
Population size changes in the host (a) and in the parasite (b) are generated for the GFG model via the parameters *b* = 1, *d* = 0.3, *δ* = 0, *β* = 0.00001, *c*_*H*_1__ = *c*_*P*_2__ = 0.05, c*H*_2_ = cP_1_ = 0 and *s* = 1. The initial conditions are *H*_1_ = *H*_2_ = 4150 and *I*_11_ = *I*_12_ = *I*_21_ = *I*_22_ = 415, so that the reference population sizes *N*_ref_ of the host and the parasite are given by 9960 and 1660, respectively, before their interaction starts at time zero. In both cases, the lower x-axes show time at the original scale of the dynamical system, whereas time is scaled by the respective values of *N*_ref_ · 1/*d* for the upper x-axes. The left y-axes denote the absolute values of the changing population size and the right y-axes denote the relative values as *ρ*(*t*) = *N*(*t*)/*N*_ref_. The number of infected alleles fluctuate between about 720 and 478.000 individuals. We evaluated allelic spectra for every second generation (on the original scale) and (c) plot the absolute number of singletons scaled by *θ, f*_20,1_(*t*)/*θ*, against time for host (dashed) and parasite (solid).

### SI 9. Distributions of singletons over time for various initial conditions

#### SI 9.1. A gene-for-gene example for 2%, 20% and 50% initially infected alleles

**Figure SI 9.1.**
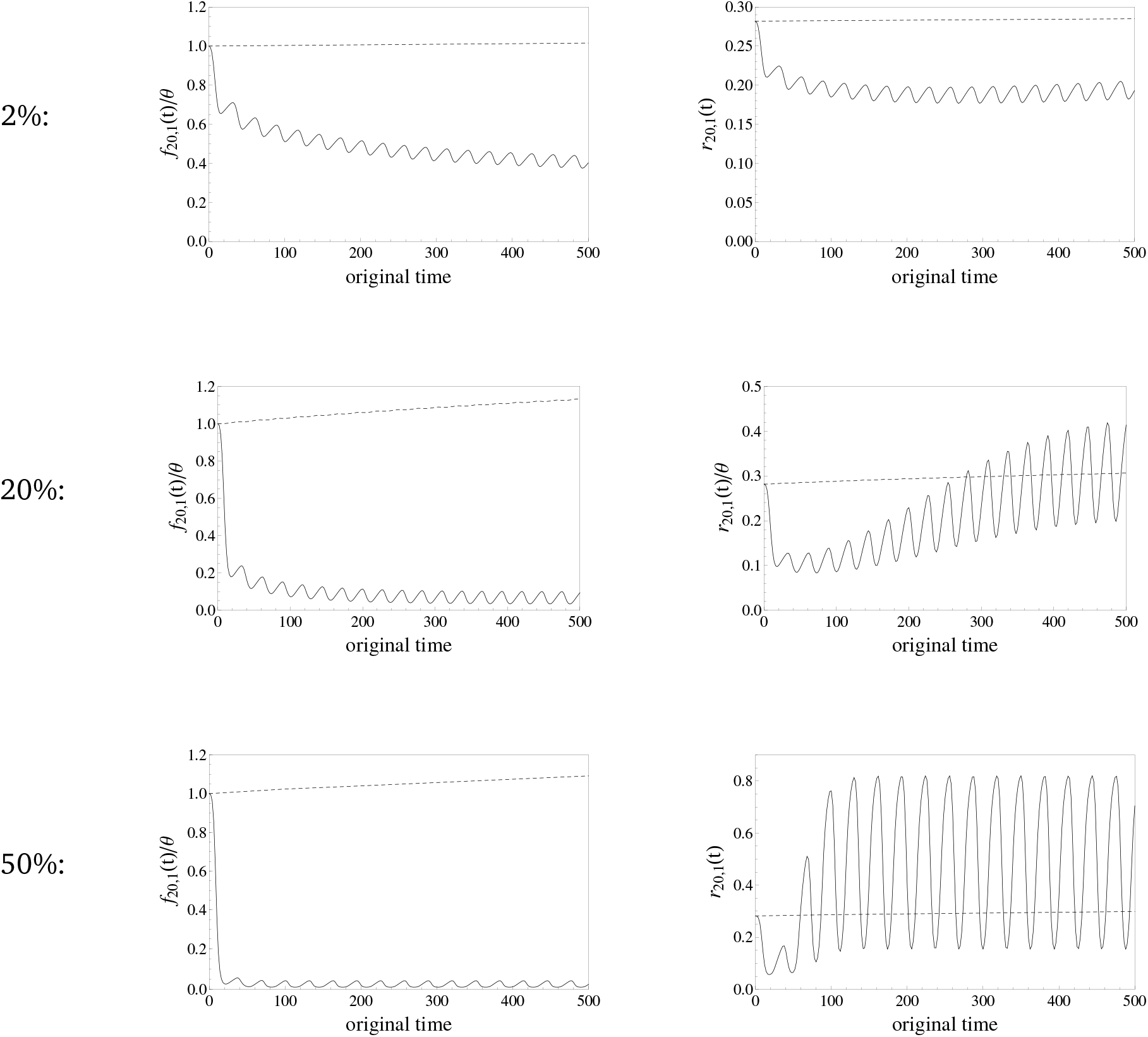
Population size changes in the host and in the parasite are generated for the GFG model via the parameters *b* = 1, *d* = 0.9, *δ* = 0.01, *c*_*H*_1__ = *c*_*P*_2__ = 0.05, *c*_*H*_2__ = *c*_*P*_1__ = 0, *s* = 1 and *β* = 0.000005 (2%) or *β* = 0.00005 (20% and 50%). The initial conditions are *H*_1_ = *H*_2_ = 49500 and *I*_11_ = *I*_12_ = *I*_21_ = *I*_22_ = 500 (2%); *H*_1_ = *H*_2_ = 4150 and *I*_11_ = *I*_12_ = *I*_21_ = *I*_22_ = 415 (20%); *H*_1_ = *H*_2_ = 3333 and *I*_11_ = *I*_12_ = *I*_21_ = *I*_22_ = 833 (50%). Therefore, the reference population sizes *N*_ref_ of the host and the parasite are, respectively, given by 101000 and 2000 (2%); 9960 and 1660 (20%); 9998 and 3332 (50%) before their interaction starts at time zero. We evaluated allelic spectra for every second generation (on the original scale) in all cases and plot the absolute number of singletons scaled by *θ*, *f*_20,1_(*t*)/θ, (left panels) and the relative number of singletons r20,1 (*t*) (right panels) against time for the host (dashed) and the parasite (solid).

#### SI 9.2. A matching-allele example for 2%, 20% and 50% initially infected alleles

**Figure SI 9.2.**
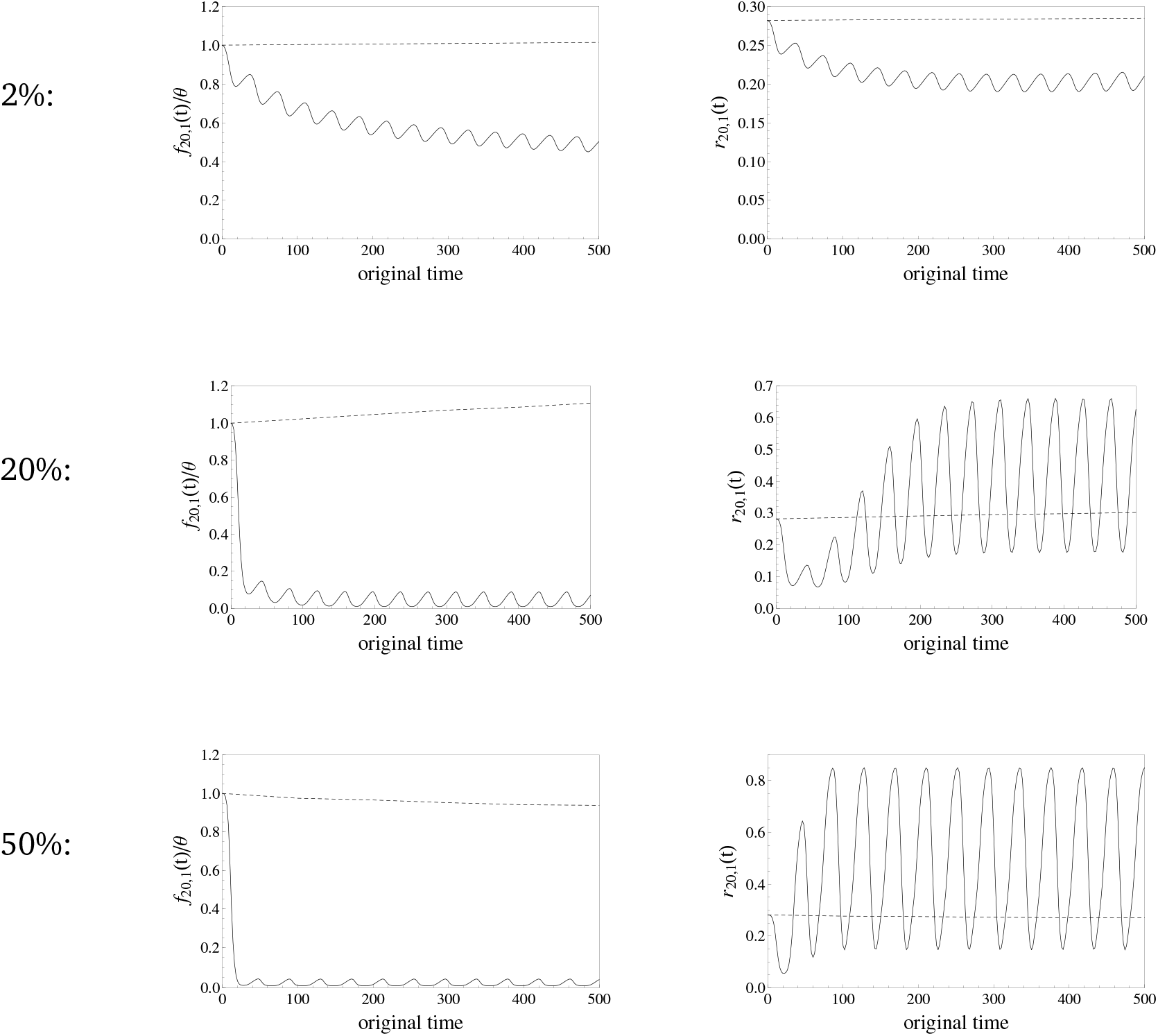
Population size changes in the host and in the parasite are generated for the MA model via the parameters *b* = 1, *d* = 0.9, *δ* = 0.01, *c_H_* = *c_P_* = 0.05, *s* = 1 and *β* = 0.0000012 (2%), *β* = 0.000012 (20%) and *β* = 0.00005 (50%). The initial conditions are *H*_1_ = *H*_2_ = 49500 and *I*_11_ = *I*_12_ = *I*_21_ = *I*_22_ = 500 (2%); *H*_1_ = *H*_2_ = 4150 and *I*_11_ = *I*_12_ = *I*_21_ = *I*_22_ = 415 (20%); *H*_1_ = *H*_2_ = 3333 and *I*_11_ = *I*_12_ = *I*_21_ = *I*_22_ = 833 (50%). Therefore, the reference population sizes *N*_ref_ of the host and the parasite are, respectively, given by 101000 and 2000 (2%); 9960 and 1660 (20%); 9998 and 3332 (50%) before their interaction starts at time zero. We evaluated allelic spectra for every second generation (on the original scale) in all cases and plot the absolute number of singletons scaled by *θ*, *f*_20,1_(*t*)/*θ*, (left panels) and the relative number of singletons *r*_20,1_(*t*) (right panels) against time for the host (dashed) and the parasite (solid).

### SI 10. The impact of multiple parasites per host and polycyclic diseases on detecting cycling population sizes for a gene-for-gene interaction

**Figure SI 10.1.**
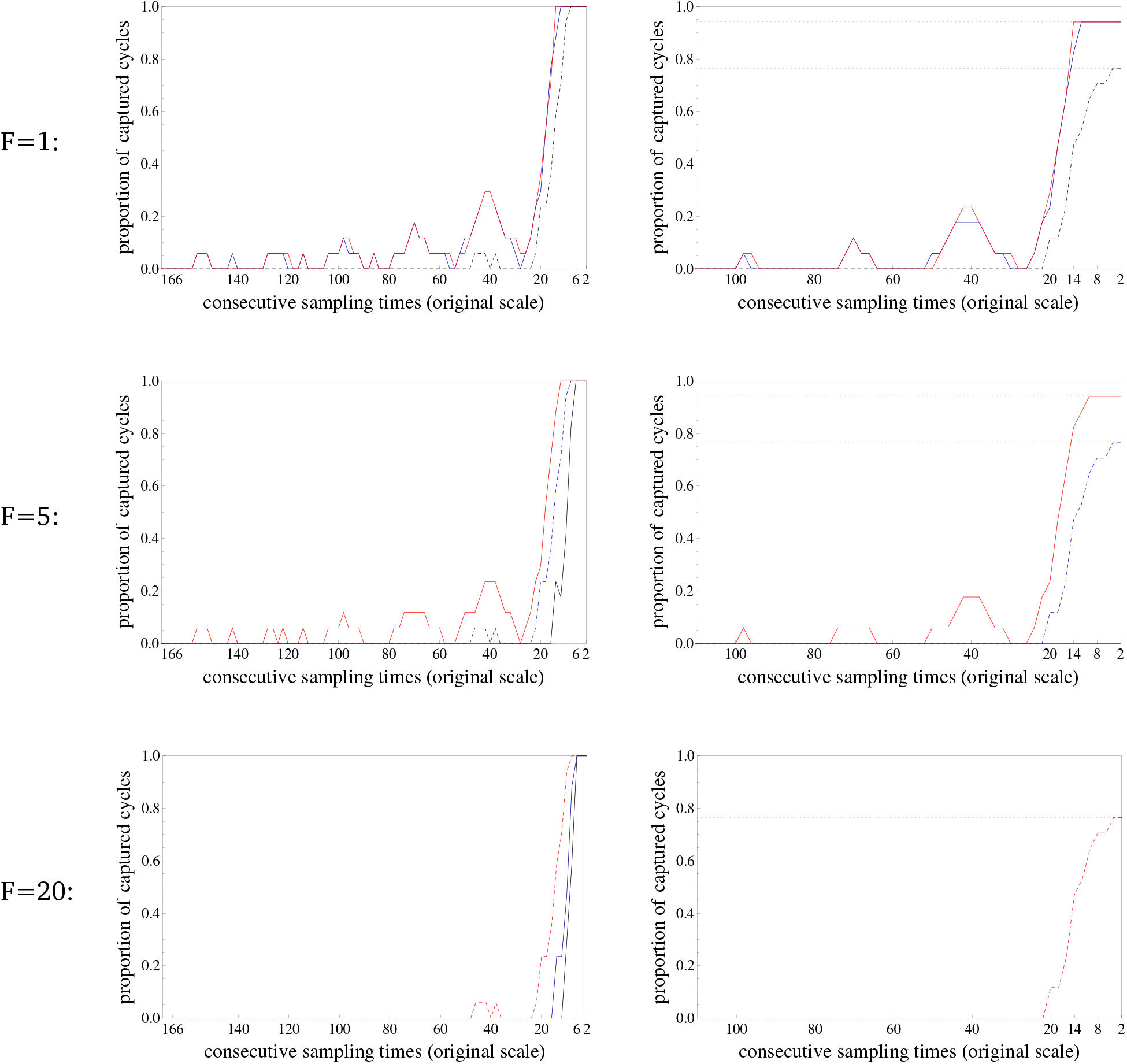

#### Description of the results

The dynamical system (1) and (2) was evaluated for the GFG model and the same parameter set as in the main example of *Results* (see Figure 3). Various values were considered for the number of parasite generations per host generation (*E* = 1, black curves; *E* = 5, blue curves; *E* = 20, red curves) and for the number of parasites per host (*F*) taking the same values as *E*. Trajectories of the population size changes of the parasite were obtained for these nine parameter combinations and respectively employed into the analytical equation of the SFS, which was evaluated at every second generation (on the original scale) over an interval of 500 time points. Based on these datasets, we evaluated Π_20_(*j* · *k*/*N*_ref_)/*θ* (see SI 4) for *j* = 2,4,6…, 166, and *k* = 0,1,2,…, up to *j* · *k* taking the largest value equal or smaller than 500. Four time points are at least required to capture a single cycle and the value *j* = 166, for instance, means that a sample is taken every 166*th* generation at times zero, 166, 332 and 498. For every *j*_0_, the times of alternating local minima and maxima of Π_20_(*j*_0_ · *k*/*N*_ref_) were obtained over the values of *k*, before the rate of change as defined by roc=(Π_20_((*m*+1)/*N*_ref_)-Π_20_(*m/N*_ref_))/Π_20_(*m/N*_ref_), *m* = 0,1,2,…, was applied until the end of the interval is reached. In all plots, the number of captured cycles are plotted (relative to the total number of cycles of the time interval) against consecutive sampling times *j*_0_. For the left-hand panels, all cycles are considered, whereas for the right-hand panels only cycles are considered, if the rate of change between two consecutive extrema deviates by at least two percent, *i.e.* roc≥ 0.02. Every cycle is captured when sampling at every sixth generation on the left-hand side whereas not all cycles can be captured in any case and not at all for all chosen values of *F < E* on the righthand side. The dashed lines illustrate cases with *E = F* giving equivalent curves. Also note that the number of captured cycles does not increase monotonically with the number of sampling times until sampling at every 20th generation in all the examples because a smaller number of sampling points (*e.g.* sampling at every 42nd generation in the first panel) can be more suitably distributed among the peaks and valleys of the considered time interval than a greater choice (*e.g.* sampling at every 30th generation in the first panel).

